# Generation and validation of a human iPSC-derived TDP-43 knockout model for ALS disease modeling

**DOI:** 10.64898/2026.04.29.720127

**Authors:** Swetha Gurumurthy, Anushka Bhargava, Nguyen PT Huynh, Thomas J. Krzystek, Fernando G. Vieira, Kyle R. Denton

**Affiliations:** ALS Therapy Development Institute, Watertown, MA, 02472

**Author notes:** These authors contributed equally to this work (Swetha Gurumurthy, Anushka Bhargava).

## Abstract

Nuclear depletion and cytoplasmic aggregation of TDP-43 occur in ∼97% of amyotrophic lateral sclerosis (ALS) cases and disrupt RNA processing through aberrant cryptic exon inclusion.

Existing cellular models rely on partial knockdown, *TARDBP* mutations, or pharmacological stress, each with limitations. Here, we generated homozygous *TARDBP*-knockout human iPSC lines using CRISPR–Cas9 genome editing and differentiated them into spinal motor neurons (MNs). Knockout MNs demonstrated ∼16-fold lower differentiation efficiency than isogenic controls but retained neuronal marker expression. TDP-43 loss induced widespread cryptic exon inclusion and depletion of STMN2, UNC13A, and G3BP1. Integration of the CUTS splice biosensor yielded up to 4.5-fold cryptic GFP induction in knockout MNs, providing a reporter-based readout of TDP-43 dysfunction. Further, we validated the cardiac glycosides digoxin and ouabain as modulators of bortezomib-induced TDP-43 pathology. This genetically defined iPSC-derived MN model provides a platform for mechanistic and therapeutic interrogation of TDP-43-driven neurodegeneration in ALS.

## Introduction

TDP-43 (TAR DNA-binding protein 43), encoded by *TARDBP*, is an essential, predominantly nuclear RNA-binding protein that regulates key aspects of RNA metabolism, including transcription, pre-mRNA splicing, mRNA stability, and transport^1^. TDP-43 is central to ALS pathogenesis, acting as both a rare genetic driver and a principal pathological substrate^2^.

Although *TARDBP* mutations account for only a minor fraction of ALS cases^3,4^, nuclear depletion and cytoplasmic aggregation of TDP-43 are observed in approximately 97% of ALS patients and represent a defining hallmark of TDP-43 proteinopathy^5^. Loss of nuclear TDP-43 is thought to be a major driver of disease through widespread disruption of TDP-43-dependent RNA processing^2,5–7^.

TDP-43 regulates pre-mRNA splicing by binding UG-rich intronic motifs and repressing the inclusion of cryptic exons^8,9^. Loss of nuclear TDP-43 leads to widespread cryptic exon inclusion, often resulting in truncated or unstable mRNAs and thereby reducing protein expression^7^. Key downstream targets of TDP-43 loss include STMN2, where loss of TDP-43 induces cryptic splicing and premature polyadenylation within intron 1, leading to near-complete loss of the full-length transcript, and UNC13A, where cryptic exon inclusion reduces normal transcript and protein abundance^10–15^. Recent studies have identified aberrant cryptic splicing in G3BP1, further linking TDP-43 dysfunction to impaired stress granule dynamics^16–20^. Together, these findings identify TDP-43 as a central regulator of neuronal RNA processing.

Human-induced pluripotent stem cell-derived motor neurons (iPSC-MNs) provide a relevant platform for studying TDP-43 function and dysfunction in ALS, recapitulating key transcriptional and functional properties of spinal MNs, the primary cell type affected in ALS, and enabling investigation of disease-relevant phenotypes in a controlled human genetic context. The development of functional TDP-43 splice reporters, including the CUTS (CFTR *UNC13A* TDP-43 loss-of-function) biosensor^21^ and the cryptic mCherry reporter^22^, enables real-time quantitative measurement of TDP-43 splicing activity in live cells and further expands the utility of these models for high-content screening.

Here, we generated and characterized a *TARDBP* knockout human iPSC model using CRISPR-Cas9 genome editing to define the consequences of complete TDP-43 loss-of-function in human iPSC-MNs. We show that TDP-43-null iPSCs differentiate into spinal motor neurons and recapitulate molecular, transcriptomic, and functional features of TDP-43 dysfunction. These include widespread cryptic exon inclusion across TDP-43 target genes, reduced expression of key downstream effectors including STMN2, UNC13A, and G3BP1, impaired neurite outgrowth, and defects in lysosomal axon transport.

We further integrated the CUTS splice reporter into *TARDBP* knockout iPSCs to enable live-cell monitoring of TDP-43 activity. Finally, we applied this model to screen small molecules reported to modulate TDP-43 mislocalization, identifying two cardiac glycosides that rescue proteasome inhibition–induced TDP-43 mislocalization. Together, this work establishes a genetically defined human iPSC-MN platform for investigating TDP-43 loss-of-function and for therapeutic screening in ALS and related TDP-43 proteinopathies.

## Results

### Generation of *TARDBP* and *STMN2* KO iPSCs

To model complete loss of TDP-43 function, we used CRISPR-Cas9 to introduce INDELs in *TARDBP* exon 1 or exon 2 (Fig. 1a) showed higher editing efficiency with the exon 2 sgRNA than with the exon 1 sgRNA (KO scores, 89 versus 23; Fig. 1b), consistent with the greater reduction in TDP-43 RNA and protein in exon 2-targeted cells^23^ (Fig. 1c–e). STMN2 protein was also reduced in exon 2 *TARDBP* KO iPSCs to a level comparable to that observed after direct targeting of STMN2 exons 2 or 3 (Fig. 1f). Following clonal isolation, selected clones showed significant reduction of TDP-43 and STMN2 mRNA and protein, increased STMN2 cryptic exon mRNA, and minimal TDP-43 protein by MSD assay (Fig. 1g–j).

**Fig 1.**
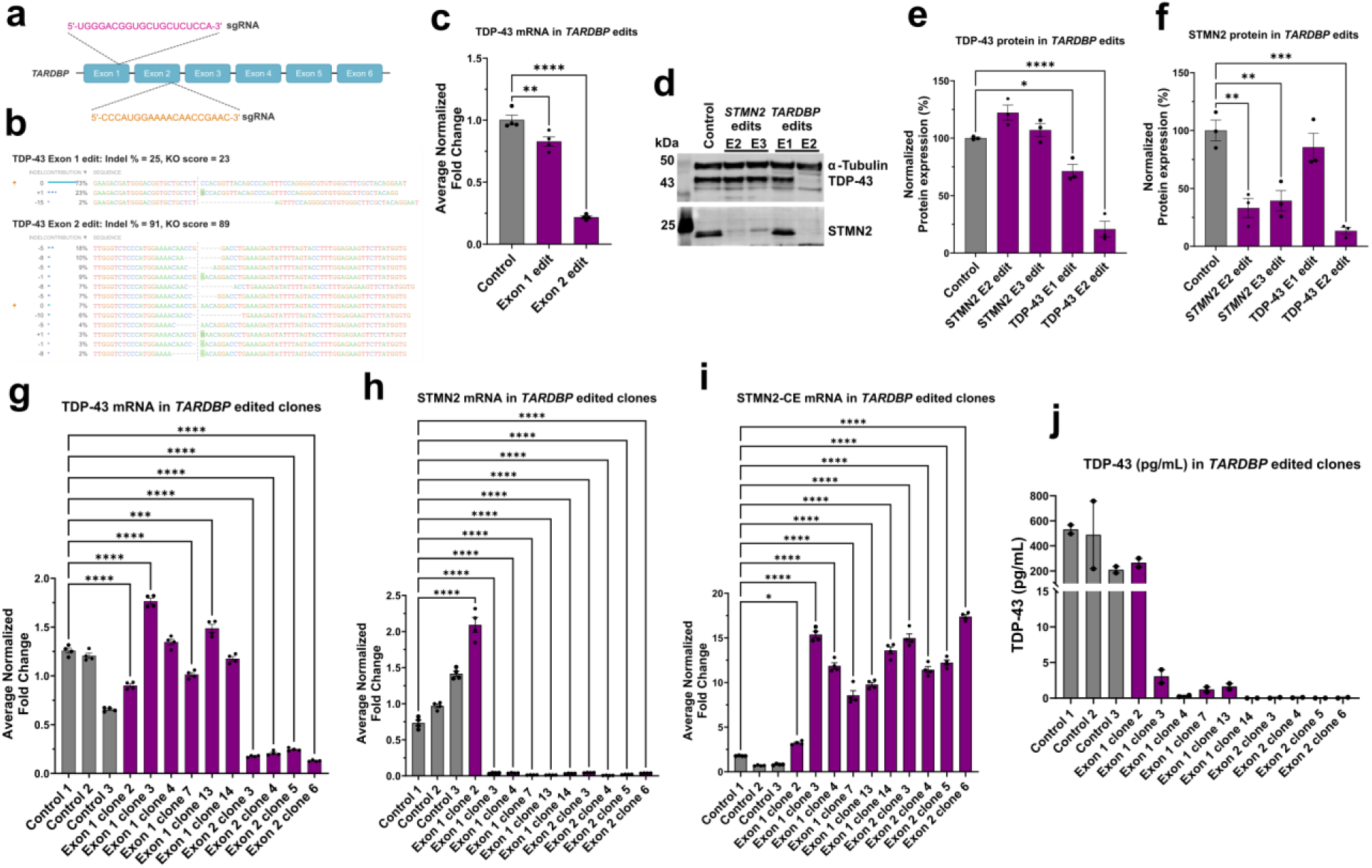
Generation and validation of homozygous *TARDBP* knockout iPSC clones. **(a)** CRISPR–Cas9 strategy used to introduce insertion/deletion (INDEL) mutations in exon 1 or exon 2 of *TARDBP*. **(b)** Editing efficiency of exon 1– and exon 2–targeting sgRNAs, as determined by ICE (Inference of CRISPR Edits) analysis. **(c)** qRT–PCR analysis of *TARDBP* mRNA expression in pooled iPSCs following sgRNA electroporation; One-way ANOVA with Dunnett’s test versus electroporation (EP)-only Control; *F*(2,9) = 171.0, *P* < 0.0001; *n* = 4. **(d)** Representative immunoblot of TDP-43, STMN2, and α-tubulin in pooled iPSC lysates following *TARDBP* knockout. **(e)** Quantification of TDP-43 protein levels in pooled edited iPSCs. Protein abundance was normalized to α-tubulin for each sample and then to EP-only Control; *F*(4,10) = 51.12, *P* < 0.0001; *n* = 3. **(f)** Quantification of STMN2 protein levels in pooled edited iPSCs. Protein abundance was normalized to α-tubulin for each sample and then to EP-only control cells; *F*(4,10) = 17.73, *P* = 0.0002; *n* = 3. **(e, f)** One-way ANOVA with Dunnett’s test versus Control. **(g)** qRT–PCR analysis of full-length *TARDBP* mRNA expression in individual iPSC clonal lines; *F*(12,39) = 1260, *P* < 0.0001; *n* = 4. **(h)** qRT–PCR analysis of full-length *STMN2* mRNA expression in individual iPSC clonal lines; *F*(12,39) = 727.6, *P* < 0.0001; *n* = 4. **(g, h)** One-way ANOVA on log-transformed data with Dunnett’s test vs Control 1. Controls 2 and 3 are shown for reference, but statistical significance is annotated only for comparisons with Control 1. **(i)** RT–PCR analysis of truncated *STMN2* cryptic exon (*STMN2*-CE) expression in individual iPSC clonal lines; *F*(12,39) = 366.4, *P* < 0.0001; *n* = 4. One-way ANOVA with Dunnett’s test versus Control 1. **(j)** Quantification of TDP-43 protein levels in individual iPSC clones using a custom Meso Scale Discovery (MSD) assay; Kruskal–Wallis with Dunn’s test versus Control 1. *H* = 23.92, *P* = 0.0208; *n* = 2. Bars represent mean ± SEM. *n* de*n*otes technical replicates per condition. \**P* < 0.05, \*\**P* < 0.01, \*\*\**P* < 0.001, \*\*\*\**P* < 0.0001.

### Generation of iPSC-MNs in the absence of TDP-43

To determine whether loss of TDP-43 is compatible with in vitro differentiation of iPSCs into spinal motor neurons, we applied an optimized dual-SMAD inhibition protocol adapted from a previously published method to generate iPSC-MNs19 (Fig. 2a). Three clones from each exon knockout were differentiated in parallel with two electroporation-only (EP-only) control lines (Supplementary Figure 1a,b). Although TDP-43 knockout lines yielded approximately 16-fold fewer motor neurons than controls, the neurons that were generated displayed comparable marker expression and culture purity (Supplementary Figure 3 and Supplementary Table 5).

**Fig 2.**
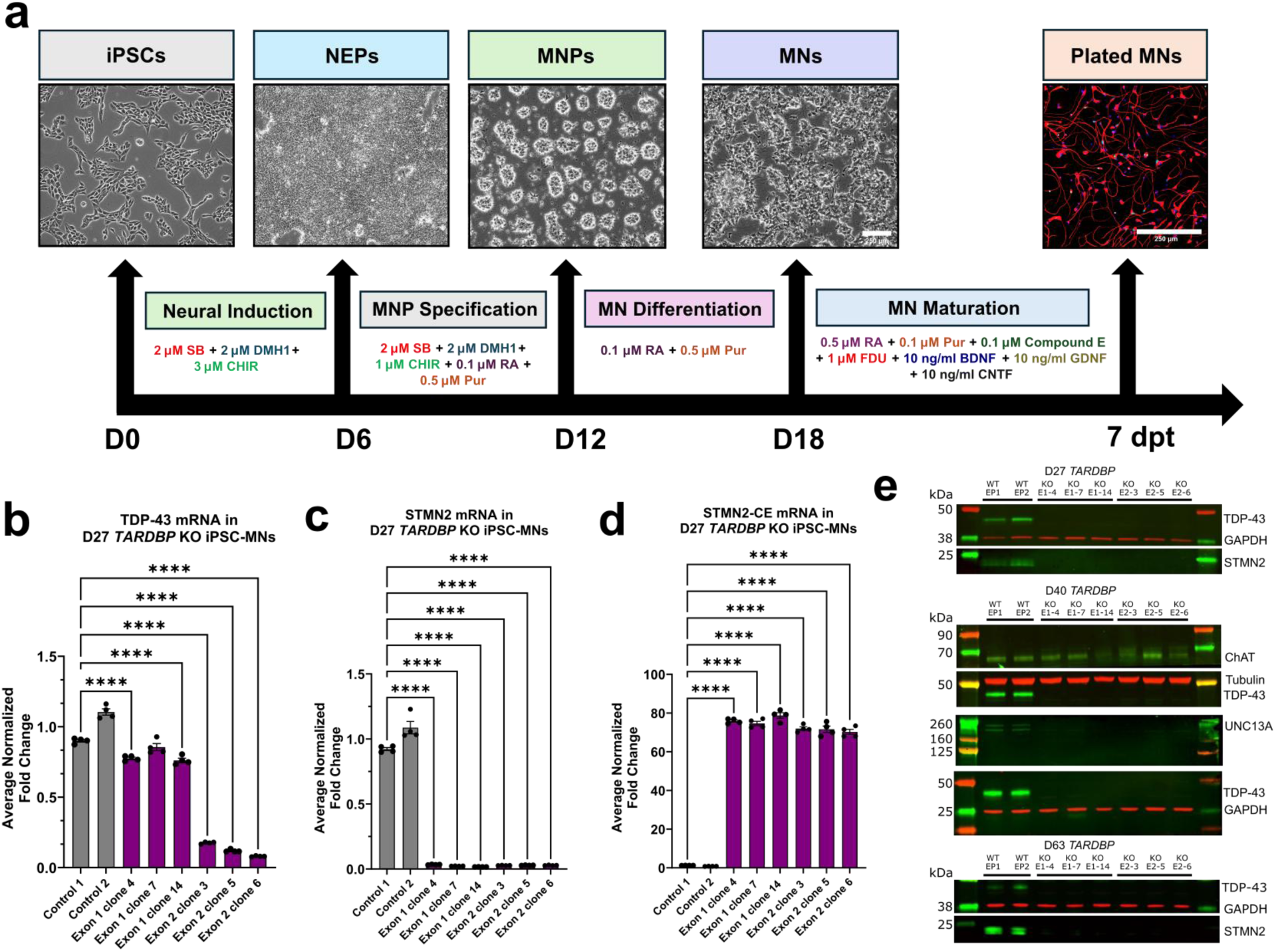
Differentiation and characterization of homozygous *TARDBP* knockout iPSC-derived motor neurons. **(a)** Schematic of the motor neuron differentiation timeline adapted from ^19^. CUTS TDP-43 WT (line 42) and CUTS TDP-43 KO (line 50) iPSCs were differentiated into neuroepithelial progenitors (NEPs) by day 6 (D6), motor neuron progenitors (MNPs) by day 12 (D12), and post-mitotic motor neurons (MNs) by day 18 (D18), followed by cryopreservation at day 19 (D19). Cryopreserved D19 MNs were subsequently thawed and matured in maturation medium. Scale bars, 250 μm. **(b)** qRT-PCR analysis of full-length *TARDBP* mRNA expression in day 27 MNs; *F*(7,24) = 816.3, *P* < 0.0001. **(c)** qRT-PCR analysis of full-length STMN2 mRNA expression in day 27 MNs; *F*(7,24) = 3402, *P* < 0.0001. **(d)** qRT-PCR analysis of truncated STMN2 cryptic exon (STMN2-CE) mRNA expression in day 27 MNs; *F*(7,24) = 11065, *P* < 0.0001. **(e)** Representative immunoblot of TDP-43 loss-of-function targets STMN2 and UNC13A at days 27, 40, and 63. **(b)** One-way ANOVA with Dunnett’s test versus Control 1. **(c, d)** One-way ANOVA on log-transformed data with Dunnett’s test versus Control 1. **(b–d)** Bars represent mean ± SEM. *n* = 4 technical replicates per condition. \**P* < 0.05, \*\**P* < 0.01, \*\*\**P* < 0.001, \*\*\*\**P* < 0.0001

### TDP-43 loss-of-function effect on splicing

To examine the effects of TDP-43 loss, we first confirmed the reduction in *TARDBP* mRNA in day 27 MNs (Fig. 2b). Next, we examined the levels of full-length and cryptic *STMN2* mRNA in day 27 MNs (Fig. 2c,d). Despite a modest reduction in *TARDBP* mRNA in the exon 1 clones, full-length *STMN2* mRNA was nearly undetectable, and STMN2-CE increased 70.3 to 78.7-fold (Fig. 2d). The loss of STMN2 was confirmed at the protein level, along with the absence of UNC13A in day 40 MNs (Fig. 2e).

To enable live-cell measurement of TDP-43 activity, we employed two recently reported splice reporter constructs^21,22^. Stable knock-in iPSC lines were generated using the SeLection by Essential-gene Exon Knock-in (SLEEK) editing strategy in which either the cryptic mCherry reporter or CUTS splice biosensor was inserted in frame at the 3’ end of the GAPDH locus of parental wild-type tdTomato or the exon 2 clone 3 TARDBP KO iPSCs, followed by isolation of clonal lines^21,22^ (Fig. 4a,b). Although the cryptic mCherry reporter did not show the expected increase in signal in TDP-43 knockout lines, the CUTS reporter performed robustly and showed up to 4.5-fold increase in nuclear cryptic GFP signal in the CUTS TDP-43 KO MNs compared to the CUTS TDP-43 WT MNs (Fig. 4c–e).

**Fig 3.**
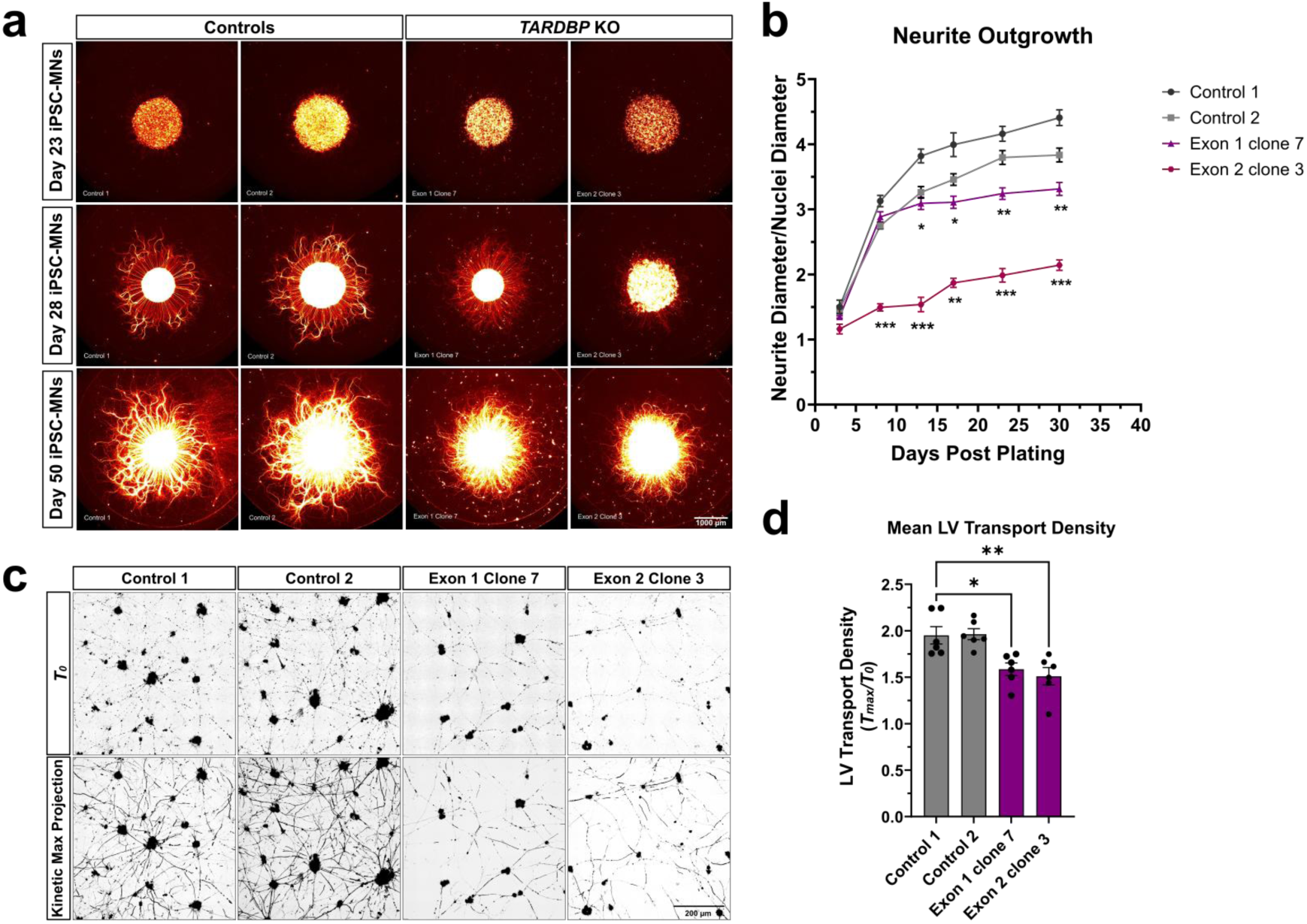
TDP-43 loss-of-function impairs neurite outgrowth and lysosome dynamics in homozygous *TARDBP* knockout iPSC-derived motor neurons. **(a)** Representative images of iPSC-MNs plated in a spot culture configuration to assess radial neurite outgrowth at days 23, 28, and 50. Scale bar, 1000 μm. **(b)** Quantification of neurite outgrowth over time, expressed as the ratio of neurite diameter to nuclear diameter. Two-way repeated measures ANOVA with Geisser–Greenhouse correction with Dunnett’s test relative to Control 1 at each time point revealed a significant time × group interaction *F*(8.765,23.37) = 28.72, *P* < 0.0001; *n* = 3 technical replicates per condition. **(c)** Representative live-cell images of day 44 MNs stained with LysoView-640. Images were acquired every second over 3 min and kinetically aligned; a kinetic maximum projection was generated to visualize lysosome transport. Scale bar, 200 μm. **(d)** Quantification of lysosome transport, calculated as the ratio of lysosome area in the maximum projection (*T_max_*) image relative to the initial (*T_0_*) image. One-way ANOVA with Dunnett’s test relative to Control 1; *F*(3, 20) = 8.82, *P* = 0.0006; *n* = 6 technical replicates per condition. **(b, d)** Bars represent mean ± SEM. \**P* < 0.05, \*\**P* < 0.01, \*\*\**P* < 0.001, \*\*\*\**P* < 0.0001.

**Fig 4:**
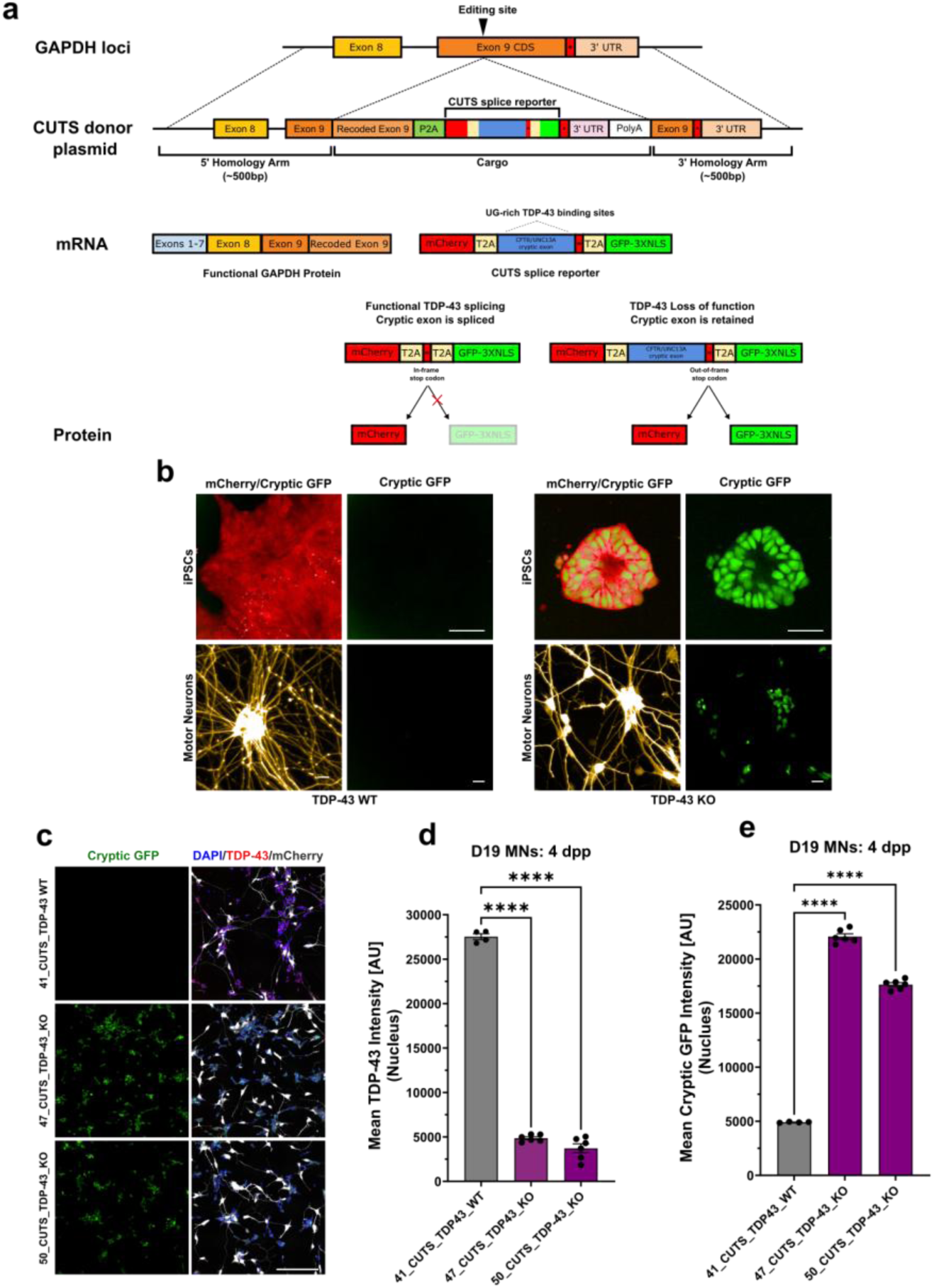
Generation of CUTS TDP-43 splice reporter iPSC lines. **(a)** Schematic of the knock-in strategy used to insert the CUTS^16^ splice reporter at the 3′ end of the *GAPDH* locus^16^. Edited clones with incorrect transgene integration are selectively eliminated due to the disruption of *GAPDH* expression. The reporter is designed to read out TDP-43-dependent suppression of cryptic exon inclusion. **(b)** Representative live fluorescence images of TDP-43 wild-type (WT) and knockout (KO) iPSCs and iPSC-MNs following CUTS reporter knock-in. Scale bars, 20 µm. **(c)** Representative immunofluorescence images of day 19 CUTS TDP-43 WT (line 41) and KO MNs (lines 47 and 50), 4dpp, showing cryptic GFP and merged images with DAPI, TDP-43, and mCherry. Scale bar, 200 µm. **(d)** Quantification of mean nuclear TDP-43 intensity in WT and KO MNs. One-way ANOVA with Dunnett’s test, *F*(2,13) = 1082, *P* < 0.0001. **(e)** Quantification of mean nuclear cryptic GFP intensity in WT and KO MNs. One-way ANOVA with Dunnett’s test, *F*(2,13) = 1586, *P* < 0.0001. **(d, e)** Bars represent mean ± SEM. WT (line 41, *n* = 4 technical replicates per condition) and KO (lines 47 and 50, *n* = 6 technical replicates per condition). \**P* < 0.05, \*\**P* < 0.01, \*\*\**P* < 0.001, \*\*\*\**P* < 0.0001.

### Functional consequences of complete TDP-43 loss-of-function

Reduced neurite outgrowth was observed in both dissociated neurons (Supplementary Figure 4a–c) and spot cultures (Fig. 3a,b). In spot cultures monitored over 50 days, *TARDBP* KO lines showed reduced neurite outgrowth relative to controls, with differences emerging after day 10 and robust by days 28 and 50 (Fig. 3a,b).

We next assessed the impact of TDP-43 loss on lysosome transport dynamics (Fig. 3c,d). Control lines showed dense, elongated lysosome trajectories, consistent with robust transport along neurites, whereas *TARDBP* KO lines exhibited fewer and shorter trajectories, indicating reduced lysosome motility. Quantification of lysosome transport density confirmed a reduction in both KO clones relative to controls (Fig. 3d), with mean transport density reduced from approximately 2.0 in controls to ∼1.5 in *TARDBP* KO lines. Together, these findings indicate that TDP-43 loss impairs endolysosomal motility in iPSC-derived motor neurons^24^.

### Long-read RNA-Seq

To evaluate the transcriptome-wide consequences of TDP-43 loss-of-function, we first examined a panel of known TDP-43-regulated cryptic exon transcripts by qRT-PCR (Supplementary Figure 2a–o). Of the 11 cryptic exon-containing transcripts that were evaluated, 9 were significantly elevated in both TDP-43 knockout (KO) lines relative to wild-type (WT) controls.

To obtain an unbiased assessment of TDP-43-dependent changes in gene expression and splicing, we performed long-read RNA sequencing on the PacBio platform to evaluate differential gene expression (DGE) and differential transcript usage (DTU) in motor neurons differentiated from wild-type (WT) and knockout (KO) lines (*n* = 2). Differential expression analysis identified 475 upregulated genes and 661 downregulated genes in KO relative to WT motor neurons (adjusted *P* ≤ 0.05). Among these, *TARDBP* expression was reduced by nearly fourfold in KO samples compared with WT controls (log2FC = −1.79, adjusted *P* ≤ 0.05; Fig. 5a and Supplementary Table 4).

**Fig 5.**
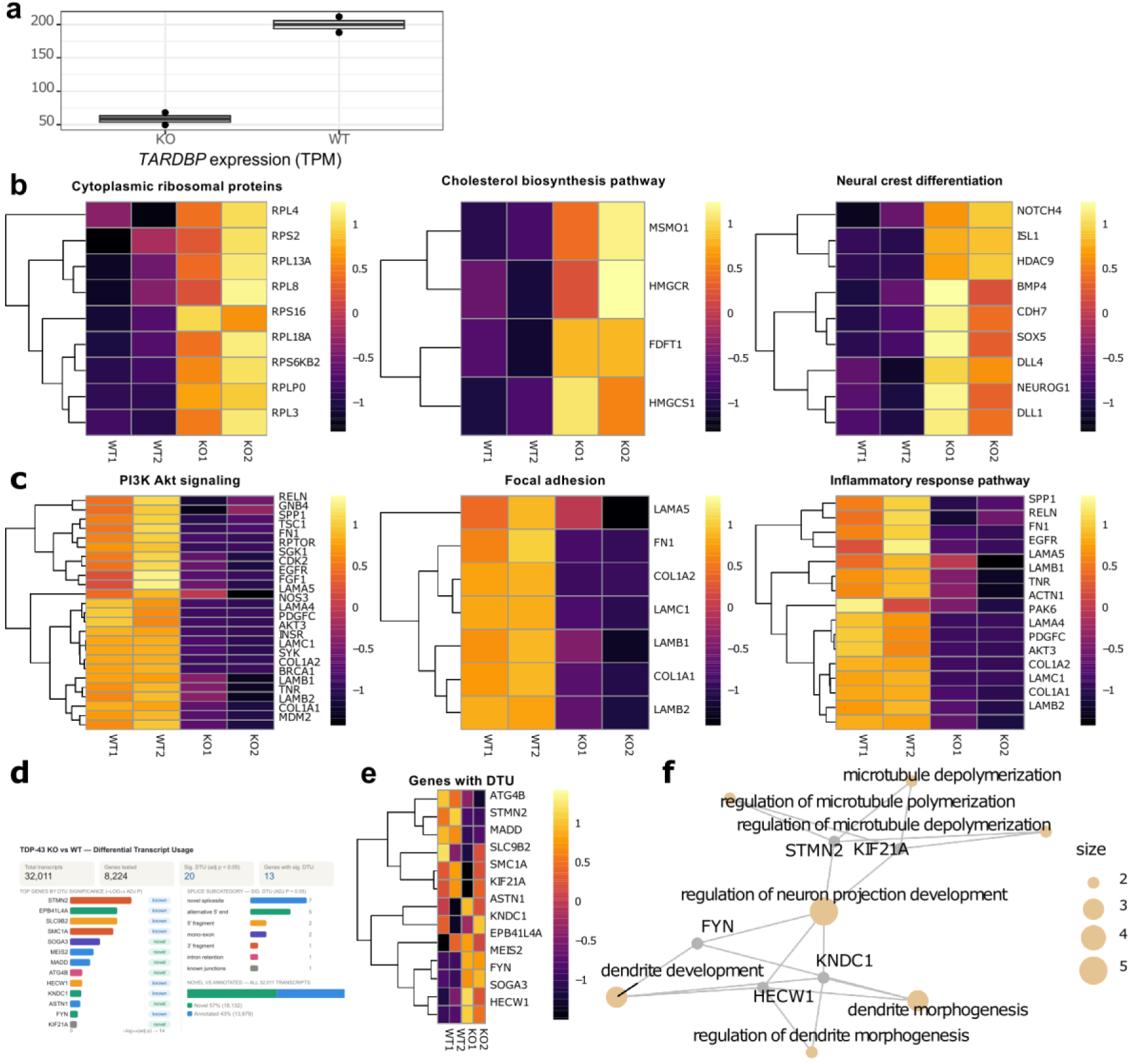
Long-read RNA-seq reveals pathway-level changes and differential transcript usage in TDP-43 knockout iPSC-derived motor neurons. **(a)** *TARDBP* expression in TDP-43 WT and KO iPSC-MNs (transcripts per million, TPM). *TARDBP* is significantly reduced in KO relative to WT (adjusted *P* ≤ 0.05). **(b, c)** Heatmaps of z-score-normalized expression for genes within selected significantly enriched gene sets in TDP-43 KO versus WT iPSC-MNs. **(b)** Pathways upregulated in KO MNs include cytoplasmic ribosomal proteins, cholesterol biosynthesis, and neural crest differentiation. **(c)** Pathways downregulated in KO MNs include PI3K-Akt signaling, focal adhesion, and inflammatory response. **(d)** Differential transcript usage (DTU) analysis across 32,011 transcripts in 8,224 genes. Shown are the top 13 genes by DTU significance with transcript status (left), splice-subcategory distribution among significant DTU events (upper right), and novel versus annotated transcript composition across all tested transcripts (lower right). **(e)** Heatmap of z-score-normalized transcript-level expression for the 13 genes with significant DTU, illustrating isoform-level shifts not reflected in total gene-level expression changes. **(f)** Network illustrates the gene ontology (GO) terms among genes with significant DTU. *n* = 2 technical replicates per genotype.

Interestingly, these data suggest that TDP-43 loss-of-function may impair motor neuron maturation, as reflected by increased expression of pathways associated with neural stemness, including neural crest differentiation, cholesterol metabolism, and ribosomal protein gene programs (Fig. 5b). In contrast, pathways linked to neuronal survival, including PI3K signaling and inflammatory response, as well as pathways related to neuronal structure, such as focal adhesion, were downregulated in TDP-43 KO motor neurons (Fig. 5c).

Given the established role of TDP-43 in isoform-specific splicing regulation, we next assessed differential transcript usage between KO and WT samples. We identified 20 isoforms from 13 genes with significant DTU (Fig. 5d). Notably, these genes did not show a consistent pattern of differential expression, with only four genes, MADD, *STMN2*, *SOGA3,* and *ATG4B,* also exhibiting significant changes at the gene level (adjusted *P* ≤ 0.05; Fig. 5e). These findings highlight the importance of transcript-level analysis in addition to gene-level analysis.

Consistent with its established role as a downstream target of TDP-43, *STMN2* exhibited altered isoform usage in TDP-43 loss-of-function motor neurons (Fig. 5d). More broadly, genes exhibiting significant DTU were predominantly associated with neuronal structural maturation.

For example, *STMN2* and *KIF21A* are key regulators of microtubule dynamics and neuronal projection, whereas *KNDC1*, *HECW1,* and *FYN* play central roles in dendrite morphogenesis and neuronal development pathways (Fig. 5f).

Taken together, these results suggest that TDP-43 loss-of-function drives a multilayered transcriptomic shift in human motor neurons. At the gene level, TDP-43 deficiency alters broad transcriptional programs consistent with impaired motor neuron maturation and survival. At the transcript level, it disrupts precise isoform usage required for neuronal structural development and function, highlighting complementary mechanisms through which TDP-43 loss contributes to motor neuron dysfunction.

### TDP-43 loss-of-function reduces STMN2 and G3BP1 protein expression in iPSC-derived motor neurons

To assess downstream protein-level consequences of TDP-43 loss-of-function, we performed quantitative immunocytochemistry in CUTS TDP-43 WT (line 42) and KO (line 50) iPSC-derived MNs. We focused on STMN2, an established downstream target of TDP-43, and G3BP1, a stress-granule scaffold linked to TDP-43-dependent RNA regulation^10,11,13,16–18^.

STMN2 was robustly detected in CUTS TDP-43 WT MNs but was markedly reduced in KO MNs, with mean cytoplasmic STMN2 intensity decreased by approximately 74% relative to WT (Fig. 6c,d). G3BP1 was likewise reduced in KO MNs, with baseline cytoplasmic G3BP1 intensity decreased by approximately 38% relative to WT (Fig. 7b). Together, these findings are consistent with previous studies and show that TDP-43 loss in this model disrupts protein pathways linked to neuronal maintenance and stress responses in MNs.

**Fig 6.**
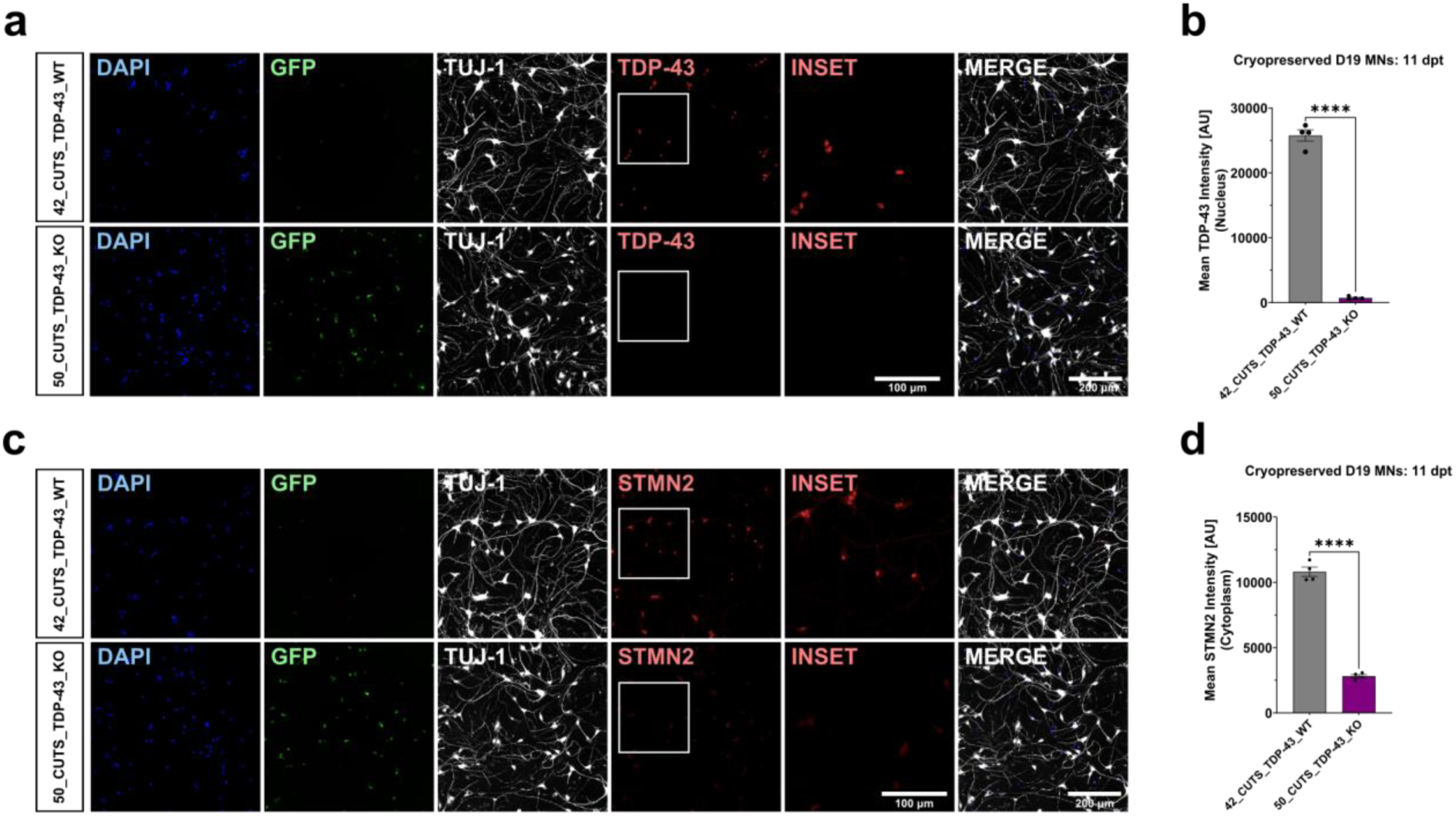
TDP-43 knockout reduces STMN2 protein expression in CUTS TDP-43 iPSC-derived motor neurons. **(a)** Representative immunofluorescence images of cryopreserved D19 CUTS TDP-43 WT (line 42) and TDP-43 KO (line 50) MNs, fixed at 11 days post-thaw, showing DAPI, cryptic GFP, TUJ-1, TDP-43, and Merge. Insets display a cropped view of the indicated TDP-43 region. **(b)** Quantification of mean nuclear TDP-43 intensity in WT and KO MNs. Two-tailed unpaired Welch’s *t*-test; *t*(3.106)=28.24, *P* < 0.0001. **(c)** Representative immunofluorescence images of cryopreserved D19 WT (line 42) and KO (line 50) MNs, fixed at 11 days post-thaw, showing DAPI, cryptic GFP, TUJ-1, STMN2, and Merge. Insets display a cropped view of the indicated STMN2 region. **(d)** Quantification of mean cytoplasmic STMN2 intensity in WT and KO MNs. Unpaired two-tailed *t*-test; *t*(6)=20.87, *P* < 0.0001. **(a, c)** Scale bars, 200 µm; insets, 100 µm. **(b, d)** Bars represent mean ± SEM. *n* = 4 technical replicates per condition. \**P* < 0.05, \*\**P* < 0.01, \*\*\**P* < 0.001, \*\*\*\**P* < 0.0001.

**Fig 7.**
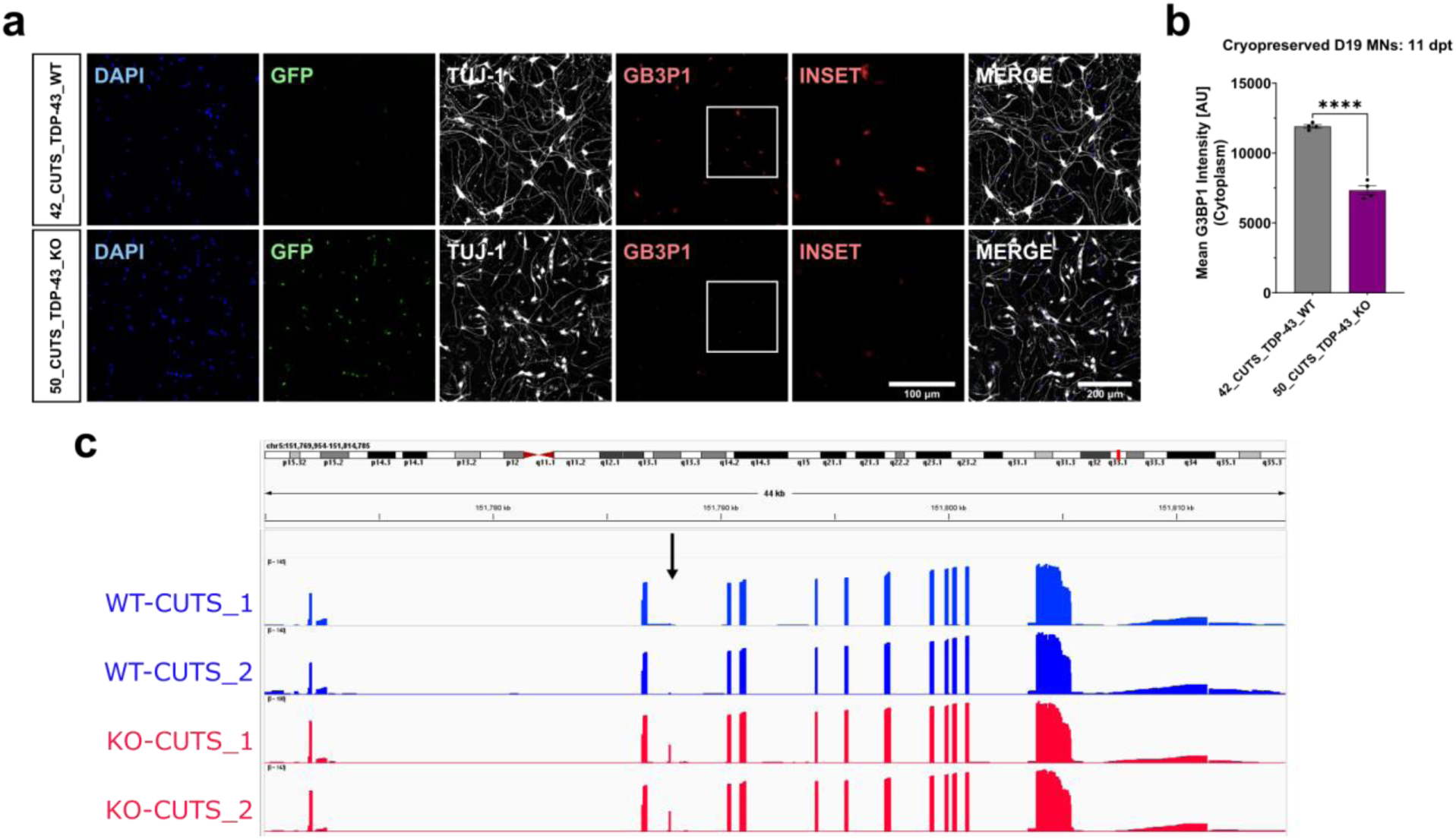
TDP-43 knockout disrupts G3BP1 expression and transcript regulation in CUTS TDP-43 iPSC-derived motor neurons. **(a)** Representative immunofluorescence images of cryopreserved D19 CUTS TDP-43 WT (line 42) and TDP-43 KO (line 50) MNs, fixed at 11 days post-thaw, showing DAPI, cryptic GFP, TUJ-1, G3BP1, and Merge. Insets show a cropped view of the indicated G3BP1 region. Scale bar, 200 µm; insets, 100 µm. **(b)** Quantification of mean cytoplasmic G3BP1 intensity in WT and KO MNs. Bars represent mean ± SEM. *n* = 4 technical replicates per condition. Two-tailed unpaired *t*-test; *t*(6) = 13.79, *P* < 0.0001. **(c)** Integrative Genomics Viewer (IGV) tracks of long read RNA-Seq coverage at the *G3BP1* locus in WT (blue) and TDP-43 KO (red) CUTS MNs. Arrow indicates a cryptic exon present in KO but absent in WT.

### Acute proteasome inhibition induces TDP-43 mislocalization in cryopreserved CUTS TDP-43 wild-type motor neurons

To assess the utility of CUTS motor neurons for small-molecule testing, we used acute proteasome inhibition to induce TDP-43 mislocalization. Treatment with either bortezomib (BTZ) or MG132 for 24 h induced TDP-43 mislocalization.

BTZ (1.1–200 nM, 24 h; IC50 for the TDP-43 mislocalization ratio, 28.26 nM) induced dose-dependent TDP-43 mislocalization in cryopreserved day 19 CUTS TDP-43 WT MNs treated at 7 days post-thaw (dpt) compared to DMSO vehicle (Fig. 8). Mean nuclear TDP-43 intensity was significantly reduced, whereas mean cytoplasmic TDP-43 intensity was significantly increased at the higher end of the dose range, resulting in a decreased nuclear-to-cytoplasmic TDP-43 ratio consistent with TDP-43 mislocalization (Fig. 8a–c,f–h). By contrast, 24 h BTZ treatment did not alter nuclear cryptic GFP reporter signal under this acute proteotoxic stress paradigm (Fig. 8d). BTZ also reduced the percentage of live nuclei and decreased total neurite length at higher concentrations (Fig. 8e,i).

**Fig 8.**
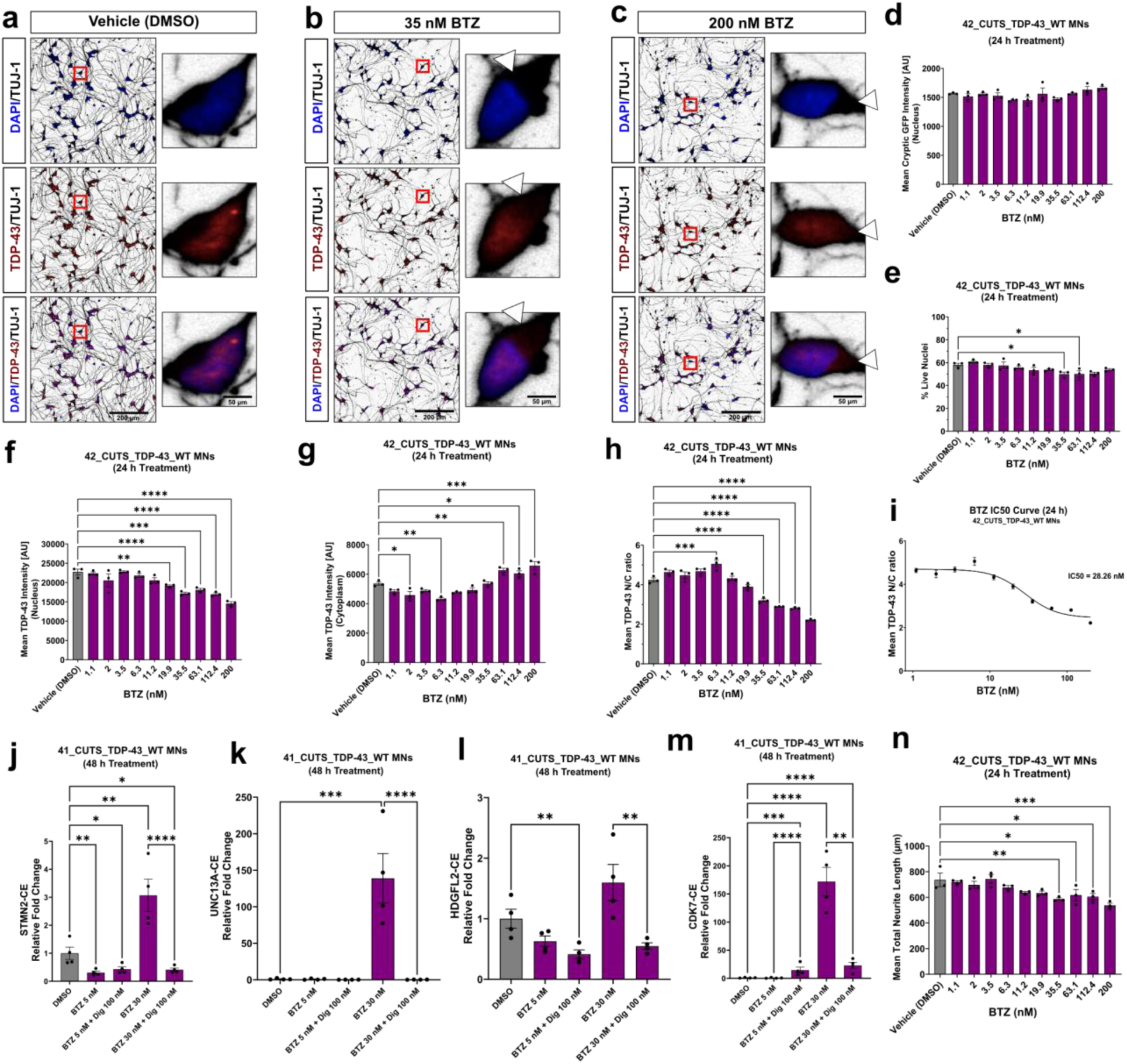
Bortezomib induces dose-dependent TDP-43 mislocalization and cryptic splicing in CUTS TDP-43 wild-type motor neurons.**(a–c)** Representative immunofluorescence images of CUTS TDP-43 WT MNs (line 42) treated for 24 h with vehicle (DMSO) (a), 35 nM BTZ (b), or 200 nM BTZ (c), showing DAPI/TUJ-1, TDP-43/TUJ-1, and merged channels. Insets highlight subcellular TDP-43 localization; white arrowheads in (b) and (c) indicate BTZ-induced TDP-43 mislocalization. **(d–h)** Quantification of mean nuclear cryptic GFP intensity (d), % live nuclei (e), nuclear TDP-43 intensity (f), cytoplasmic TDP-43 intensity (g), and TDP-43 nuclear/cytoplasmic (N/C) ratio (h) in WT MNs treated with BTZ (1.1–200 nM, 24 h). **(i)** IC50 curve for TDP-43 mislocalization (BTZ, 24 h); four-parameter variable-slope model (log[inhibitor] versus response); IC50 = 28.26 nM. **(j–m)** Relative fold change in cryptic exon expression for STMN2-CE (j; *F*(4,15) = 27.06, *P* < 0.0001), UNC13A-CE (k; *F*(4,15) = 23.55, *P* < 0.0001), HDGFL2-CE (l; *F*(4,15) = 11.69, *P* = 0.0002) and CDK7-CE (m; *F*(4,15) = 46.69, *P* < 0.0001) in CUTS TDP-43 WT MNs (line 41) treated for 48 h with 5 nM or 30 nM BTZ alone or together with 100 nM digoxin. One-way ANOVA on log-transformed data with Dunnett’s test versus Vehicle (DMSO); *n* = 4 technical replicates per condition. **(n)** Quantification of mean total neurite length (BTZ, 24 h). **(d–h, n)** One-way ANOVA with Dunnett’s test versus Vehicle (DMSO); *n* = 3 technical replicates per condition. Quantified parameters were mean nuclear cryptic GFP intensity (d; *F*(10,22) = 2.190, *P* = 0.0603), % live nuclei (e; *F*(10,22) = 4.057, *P* = 0.0030), nuclear TDP-43 intensity (f; *F*(10,22) = 16.22, *P* < 0.0001), cytoplasmic TDP-43 intensity (g; *F*(10,22) = 20.78, *P* < 0.0001), TDP-43 N/C ratio (h; *F*(10,22) = 72.73, *P* < 0.0001) and mean total neurite length (n; *F*(10,22) = 5.762, *P* = 0.0003). **(a–c)** Scale bars, 200 µm; insets, 50 µm. **(d–n)** Bars represent mean ± SEM.

To determine whether BTZ-induced TDP-43 mislocalization was accompanied by downstream mis-splicing, we performed qRT-PCR on WT CUTS motor neurons at 3 days post-plating following treatment with DMSO, 5 nM BTZ, or 30 nM BTZ (Fig. 8j–m). Treatment with 30 nM BTZ increased expression of *STMN2*-CE, *UNC13A*-CE, and *CDK7*-CE relative to DMSO-treated cells, indicating that acute proteasome inhibition induces cryptic exon inclusion in this model.

MG132 (5.6–1,000 nM, 24 h; IC50 for the TDP-43 mislocalization ratio, 270.7 nM) similarly induced dose-dependent TDP-43 mislocalization in cryopreserved day 19 CUTS TDP-43 WT motor neurons treated at 7 days post-plating relative to DMSO vehicle (Supplementary Figure 5a–k). Mean nuclear TDP-43 intensity was reduced, whereas mean cytoplasmic TDP-43 intensity was increased, resulting in a decreased nuclear-to-cytoplasmic TDP-43 ratio (Supplementary Figure 5a–c,f–h). Nuclear cryptic GFP reporter expression remained unchanged across the concentration range tested (Supplementary Figure 5d). Unlike BTZ, 24 h MG132 treatment did not significantly alter the percentage of live nuclei across the concentrations tested (Supplementary Figure 5e), whereas total neurite length was reduced at the highest concentrations (Supplementary Fig. 5k). Together, these findings show that acute proteasome inhibition by BTZ and MG132 induces dose-dependent TDP-43 mislocalization in the CUTS TDP-43 motor neuron model, with higher concentrations additionally compromising neurite integrity.

### Digoxin and ouabain rescue BTZ-induced TDP-43 proteinopathy in cryopreserved CUTS TDP-43 wild type motor neurons

Treatment of cryopreserved day 19 CUTS TDP-43 WT motor neurons with 30 nM BTZ for 48 h induced marked TDP-43 mislocalization relative to the DMSO vehicle control (Fig. 9a,b,e–h). Co-treatment with digoxin increased the TDP-43 nuclear-to-cytoplasmic intensity ratio, the percentage of live nuclei, and total neurite length, consistent with rescue of BTZ-induced TDP-43 mislocalization and neurite degeneration (Fig. 9c,i,m). By contrast, digoxin did not significantly alter mean nuclear cryptic GFP intensity relative to BTZ alone (Fig. 9k). Although treatment with 5 nM BTZ for 48 h did not significantly affect cryptic transcript levels, 30 nM BTZ significantly increased expression of STMN2-CE, UNC13A-CE, and CDK7-CE (Fig. 8j–m). Co-treatment with 100 nM digoxin reversed this BTZ-induced increase in cryptic transcript expression (Fig. 8j–m).

**Fig 9.**
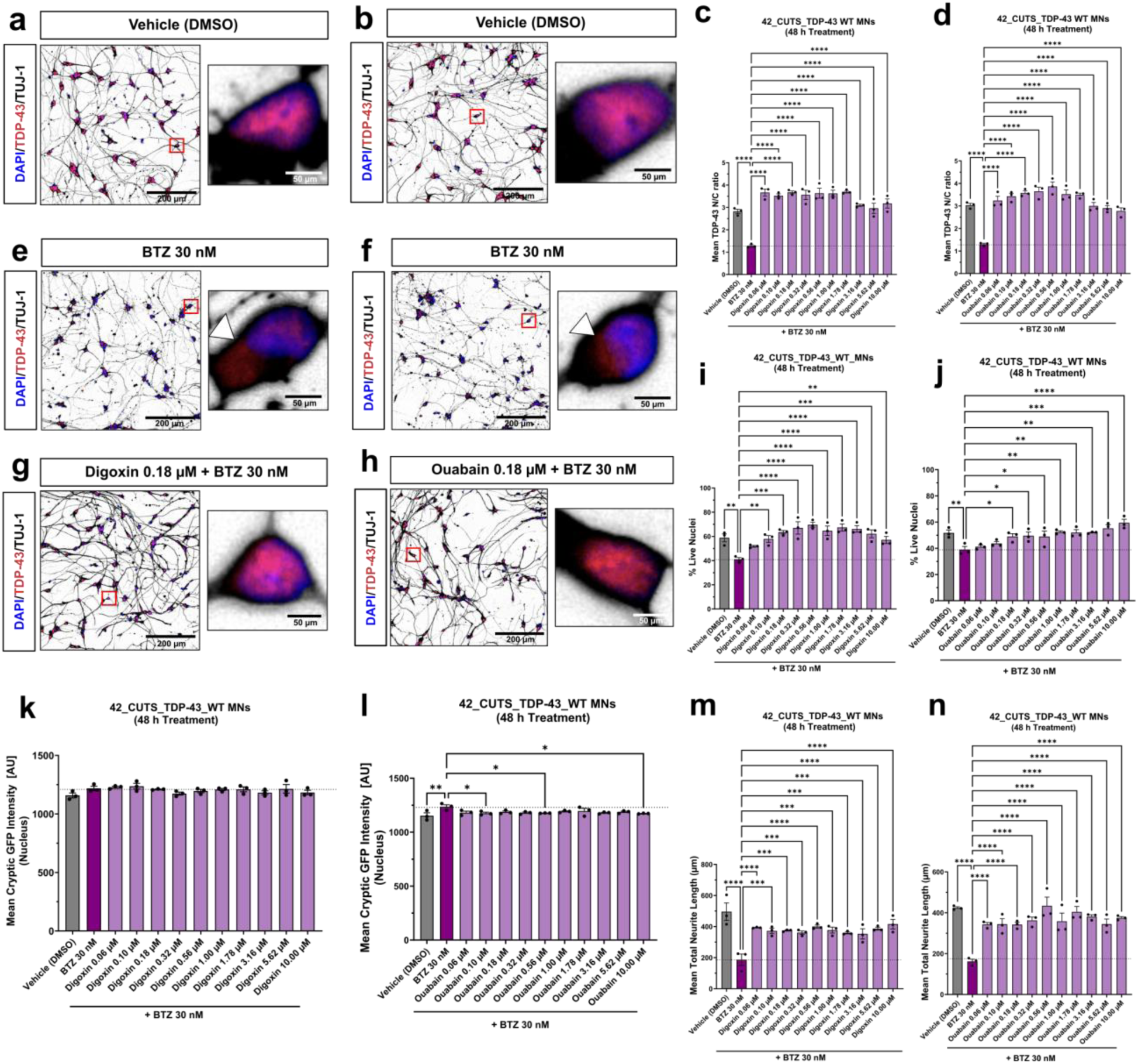
Digoxin and ouabain rescue bortezomib-induced TDP-43 pathology in CUTS TDP-43 wild-type motor neurons. **(a, b, e, f, g, h)** Representative immunofluorescence images of cryopreserved D19 CUTS TDP-43 WT MNs (line 42) treated for 48 h and fixed at 11 days post-thaw. Images show a merge of DAPI, TDP-43, and TUJ-1. Insets show cropped views of the highlighted cells. **(a, b)** DMSO vehicle controls corresponding to digoxin (a) and ouabain (b). **(c, d)** Quantification of the mean TDP-43 nucleo-cytoplasmic (N/C) ratio across digoxin (c; *F*(11, 24) = 21.66, *P* < 0.0001) and ouabain titrations (d; *F*(11, 24) = 21.93, *P* < 0.0001). **(e, f)** Representative immunofluorescence images of WT MNs treated with bortezomib (BTZ) 30 nM alone for comparison within the digoxin (e) and ouabain (f) titrations; arrowheads indicate cytoplasmic mislocalization of TDP-43. **(g, h)** Representative immunofluorescence images of WT MNs co-treated with BTZ 30 nM and digoxin 0.18 μM (g) or ouabain 0.18 μM (h); insets highlight nuclear restoration of BTZ-mislocalized TDP-43. **(i, j)** Quantification of % live nuclei across digoxin (i; *F*(11, 24) = 7.408, *P* < 0.0001) and ouabain titrations (j; *F*(11, 24) = 6.096, *P* = 0.0001). **(k, l)** Quantification of mean nuclear cryptic GFP intensity with 30 nM BTZ co-treatment across digoxin (k; *F*(11, 24) = 1.512, *P* = 0.1912) and ouabain titration (l; *F*(11, 24) = 2.026, *P* = 0.0719). **(m, n)** Quantification of mean total neurite length (μm) across digoxin titration (m; *F*(11, 24) = 7.720, *P* < 0.0001) or ouabain titration (n; *F*(11, 24) = 9.185, *P* < 0.0001). **(a, b, e, f, g, h)** Scale bars, 200 μm; insets, 50 μm. **(c, d, i, j, k–n)** Bars represent mean ± SEM; *n* = 3 technical replicates per condition. One-way ANOVA with Dunnett’s test vs BTZ 30 nM. \**P* < 0.05, \*\**P* < 0.01, \*\*\**P* < 0.001, \*\*\*\**P* < 0.0001.

Ouabain produced a similar rescue effect, significantly increasing the TDP-43 nuclear-to-cytoplasmic intensity ratio, restoring the percentage of live nuclei, and increasing total neurite length (Fig. 9d,j,n). In addition, ouabain reduced mean nuclear cryptic GFP intensity at selected concentrations relative to BTZ alone (Fig. 9l). Together, these findings identify digoxin and ouabain as robust modulators of BTZ-induced TDP-43 proteinopathy in the CUTS TDP-43 motor neuron model.

## Discussion

In this study, we generated homozygous *TARDBP* knockout iPSC lines and differentiated them into spinal motor neurons that recapitulate the molecular and functional features of TDP-43 loss-of-function. We also established the CUTS TDP-43 splice reporter in human iPSC-derived motor neurons, providing a tractable platform for monitoring TDP-43-dependent splicing dysfunction.

A central finding was that homozygous *TARDBP* knockout iPSCs retained the capacity to generate spinal motor neurons, with neuronal marker expression broadly comparable to controls through day 60. However, knockout lines produced approximately 16-fold fewer motor neurons, indicating that complete *TARDBP* loss impairs survival during differentiation, particularly at the motor neuron progenitor and early post-mitotic stages. These data suggest an early requirement for TDP-43 in efficient motor neuron production and point to reduced progenitor and/or early neuronal fitness as a major consequence of *TARDBP* loss.

Long-read RNA sequencing of *TARDBP* knockout motor neurons revealed a coordinated transcriptomic shift consistent with impaired maturation. At the gene-expression level, knockout motor neurons upregulated cytoplasmic ribosomal protein, cholesterol biosynthesis, and neural crest differentiation programs, while downregulating PI3K-Akt signaling, focal adhesion, and inflammatory response pathways (Fig. 5b,c). Retention of neural crest and early developmental markers, including NEUROG1, ISL1, BMP4, and NOTCH4, together with reduced expression of mature neuronal signaling and cell-matrix adhesion programs, suggests that TDP-43 loss arrests motor neuron differentiation in a more primitive transcriptional state. Upregulation of ribosomal protein and cholesterol biosynthesis genes is also consistent with elevated translational and biosynthetic activity characteristic of less differentiated cells.

At the transcript level, differential transcript usage analysis identified 20 isoforms across 13 genes with significant isoform shifts in *TARDBP* knockout motor neurons (Fig. 5d). Notably, only four of these genes, MADD, STMN2, SOGA3, and ATG4B, also exhibited differential expression, indicating that isoform-resolution profiling captures TDP-43-dependent regulation not apparent at the gene level. Differential transcript usage (DTU) genes were enriched for neuronal projection development, dendrite morphogenesis, and microtubule dynamics (Fig. 5f), consistent with a role for TDP-43 in coordinating the isoform repertoire required for cytoskeletal remodeling during motor neuron maturation. STMN2, a well-established downstream target of TDP-43-dependent splicing repression, was among the top differential transcript usage genes, providing internal validation.

Functionally, *TARDBP* knockout motor neurons showed impaired neurite outgrowth and reduced lysosome transport in the absence of exogenous stress, indicating that TDP-43 loss alone is sufficient to induce disease-relevant motor neuron dysfunction. Reduced lysosome motility is also consistent with broader defects in axonal trafficking linked to ALS^24^. Integration of the CUTS reporter into exon 2 *TARDBP* KO iPSCs enabled a quantitative single-cell readout of TDP-43-dependent splicing dysfunction. KO MNs showed an approximately 4.5-fold increase in nuclear cryptic GFP signal relative to wild-type MNs, confirming reporter sensitivity and establishing a scalable readout for high-content screening. Quantitative immunofluorescence further revealed reduced STMN2 and G3BP1 protein levels in *TARDBP* KO MNs (Figs. 6 and 7), linking TDP-43 loss to disruption of both axonal maintenance and stress granule competence.

Given the link between proteasome dysfunction and TDP-43 pathology in ALS, we induced proteotoxic stress in wild-type CUTS TDP-43 MNs using bortezomib and MG132 and assessed the effects of the cardiac glycosides digoxin and ouabain^25–27^. Under bortezomib-induced stress, both compounds improved the TDP-43 nuclear-to-cytoplasmic ratio, increased viability, and preserved neurite integrity across the tested concentration range. However, digoxin did not significantly alter the nuclear CUTS cryptic GFP reporter signal, whereas ouabain reduced this signal relative to bortezomib alone. These findings support a neuroprotective effect of cardiac glycosides in this context while suggesting differential effects on TDP-43-associated cryptic splicing.

This study has several limitations. Homozygous *TARDBP* KO represents a more severe loss-of-function state than the partial and heterogeneous nuclear depletion of TDP-43 observed in patient tissue, and future studies should determine whether graded TDP-43 reduction produces proportionate molecular and functional phenotypes. Although KO MNs remained viable through day 60, longer-term culture will be needed to assess progressive degeneration. In addition, the long-read RNA-sequencing analysis was performed with two biological replicates per genotype, limiting power to detect subtler transcript-level changes. Finally, testing whether restoration of downstream targets, including STMN2, UNC13A, and G3BP1, rescues key phenotypes will further strengthen this platform for therapeutic discovery.

Together, we generated and validated a homozygous *TARDBP* KO human iPSC MN model that combines a live-cell CUTS TDP-43 splice reporter with immunocytochemical analysis of TDP-43, STMN2, and G3BP1. In this platform, we identified cardiac glycosides as modulators of TDP-43 proteinopathy and cryptic exon activity. This model provides a framework for dissecting TDP-43-driven neurodegeneration and for evaluating therapeutic strategies aimed at restoring TDP-43-dependent RNA processing or mitigating its downstream pathological consequences in ALS-related TDP-43 proteinopathies.

## Methods

### Human induced pluripotent stem cell culture

iPSCs were maintained in mTeSR Plus (STEMCELL Technologies, 100-0276) on Matrigel-coated plates (Corning, 354277) and passaged at 70% confluency using Accutase (STEMCELL Technologies, 07920). Following dissociation, cells were centrifuged at 200 × g for 5 min and resuspended in mTeSR Plus supplemented with 10 μM Y-27632 (SelleckChem, S1049). All iPSC lines were passaged at least twice post-thawing prior to differentiation.

### sgRNA Guide Design

sgRNAs targeting exon 1 or exon 2 of *TARDBP* were designed using Benchling’s CRISPR guide RNA design tool and selected based on predicted high on-target efficiency and low off-target activity. Synthetic chemically modified sgRNAs were purchased from Synthego, resuspended in TE buffer to a final concentration of 100 μM, and stored at −80 °C until use. The sgRNA sequences were as follows: *TARDBP* exon 1, 5’-CCCAUGGAAAACAACCGAAC-3’; *TARDBP* exon 2, 5’-UGGGACGGUGCUGCUCUCCA-3’; STMN2 exon 2 forward, 5’-AUCAACAUCUAUACUUACGA-3’; STMN2 exon 2 reverse, 5’-AGAGCAGAUCAGUGACAGCA-3’; STMN2 exon 3 forward, 5’-GUUCGUGGGGCUUCUGAGAU-3’; and STMN2 exon 3 reverse, 5’-GAAGAAAGACCUGUCCCUGG-3’. The SLEEK GAPDH guide sequence was 5’-TCTAGGTATGACAACGAATTGUUUUAGAGCUAGAAAUAGCAAGUUAAAAUAAGGCUAGUCCGUUAUCAACUUGAAAAAGUGGCACCGAGUCGGUGCUUUU-3’.

### Genome editing

*TARDBP* knockout iPSCs were generated from the previously published VACHT-tdTomato spinal motor neuron reporter line^28^. To introduce double-stranded DNA breaks in either exon 1 or exon 2 of *TARDBP*, iPSCs were electroporated with the ribonucleoprotein (RNP) complexes consisting of Alt-R SpCas9 Nuclease V3 (IDT, 1081059) and each of the different sgRNA guides (Synthego, sgRNA EZ kit) at a 1 to 3 molar ratio, respectively. Briefly, iPSCs were dissociated using Accutase for 3 minutes at 37°C in a hypoxic incubator. Cells were then collected in 10 mL PBS and centrifuged at 200 x g for 5 minutes. For each electroporation, at least 1 × 10^6^ cells were resuspended in Neon Buffer R. RNP complexes were assembled by incubating Cas9 protein with both sgRNAs for 20 min at room temperature, and then added to the dissociated cells in Neon Buffer R. Electroporation was performed using the Neon Transfection System (Thermo Fisher Scientific) with the following settings: 1100 V, 20 ms, 1 pulse.

### Clonal isolation and cryopreservation

Clonal lines were generated by plating 1 × 10^5^ cells in a 15-cm dish and manually picking individual colonies. Cells were banked at 1 ×10^6^ cells or 5 ×10^5^ cells per vial in 10% DMSO and 90% filter-sterilized KnockOut Serum Replacement (Thermo Fisher Scientific, 10828028).

### DNA isolation, PCR, and Sanger sequencing

Two days after plating, individual clones were washed with DPBS and lysed in QuickExtract DNA Extraction Solution (Lucigen, QE09050). Lysates were mixed by pipetting ten times and stored at-80 °C until analysis. For PCR, 1 µL lysate was used as template in reactions containing HotStart polymerase and 10 µM primers flanking the target site. PCR products were analyzed by Sanger sequencing (GENEWIZ, South Plainfield, NJ, USA). Editing efficiency and knockout genotype were determined using Synthego ICE Analysis v3.0.

### Motor neuron differentiation

Motor neuron differentiation was performed using a published protocol adapted from^29^. Neural differentiation medium (NDM) consisted of a 1:1 mixture of DMEM/F12 and Neurobasal medium supplemented with N2, B27, penicillin–streptomycin, GlutaMAX, MEM non-essential amino acids, and 0.1 mM ascorbic acid. For neuroepithelial progenitor induction, *TARDBP* WT and KO iPSCs were dissociated with Accutase and plated on Matrigel-coated plates at 30,000 cells/cm². On day 0, iPSCs were transitioned to NDM supplemented with 3 µM CHIR99021, 2 µM DMH-1, and 2 µM SB431542, with daily medium changes for 6 d. On day 7, neuroepithelial progenitors were dissociated and replated at 300,000 cells/cm² in NDM supplemented with 0.1 µM retinoic acid, 0.5 µM purmorphamine, 1 µM CHIR99021, 2 µM DMH-1, and 2 µM SB431542 to generate motor neuron progenitors (MNPs). On day 13, MNPs were dissociated and cultured in NDM supplemented with 0.5 µM retinoic acid and 0.1 µM purmorphamine, with daily medium changes for 6 d, generating post-mitotic MNs by day 19. For neuronal maturation, day 19 MNs were seeded in motor neuron plating medium consisting of NDM supplemented with 0.5 µM retinoic acid, 0.1 µM purmorphamine, 0.1 µM Compound E, 10 µM Y-27632 (ROCK inhibitor), 5 µM 5-fluoro-2′-deoxyuridine (FUdR), and 10 ng/mL each of brain-derived neurotrophic factor (BDNF), ciliary neurotrophic factor (CNTF), and glial cell line-derived neurotrophic factor (GDNF). After 48 h, half-medium changes were performed every other day using the same medium lacking Y-27632 2HCl and FUdR.

### Neuronal spot cultures

The spot culture method was adapted from a previously described protocol^12^. iPSC-MNs were plated in 96-well plates pre-coated with poly-L-ornithine (PLO;100 µg/mL) and laminin (10 µg/mL). For each well, 1 µL of cell suspension at 1 × 10^5^ cells/µL was dispensed into the center of the well. Plates were incubated in a humidified chamber at 37 °C for 10 min to allow cell attachment, after which 100 µL of motor neuron plating medium was gently added without disturbing the adherent spot. Half-medium changes were performed every 3 d.

### Quantification of TDP-43 by R-PLEX immunoassay

TDP-43 was quantified using the R-PLEX Human TDP-43 Assay (Meso Scale Discovery, Cat. No. K151AMQR) on a MESO QuickPlex SQ 120MM reader, following the manufacturer’s instructions. An eight-point standard curve was prepared from the 20× calibrator stock in Assay Diluent 12 by diluting 15 µl calibrator into 285 µl diluent for the first standard, followed by 1:4 serial dilutions. Cell lysates were prepared in RIPA buffer supplemented with 2× protease inhibitors. Samples were serially diluted to target concentrations in PBS containing 2× protease inhibitors. MSD GOLD Small Spot Streptavidin plates were coated with 25 µL per well of biotinylated capture antibody (200 µL capture antibody in 3.3 mL Diluent 12) and incubated for 1 h at room temperature with shaking at 700–800 rpm. Plates were washed three times with 150 µL per well of PBS containing 0.05% Tween-20, after which 25 µl Assay Diluent and 25 µl calibrator or sample were added to each well. Plates were sealed and incubated for 1 h at room temperature with shaking. After a second wash, 50 µl per well of 1× SULFO-TAG-conjugated detection antibody (60 µL detection antibody in 6 mL Diluent 11) was added to each well and incubated for 1 h at room temperature with shaking. Following a final wash, 150 µL of MSD GOLD Read Buffer B was added to each well, and plates were read immediately on the SQ 120 instrument. Data were analyzed using Discovery Workbench v4.0. Standard curves were fitted using a four-parameter logistic model with 1/y² weighting. The lower limit of detection (LLOD) was defined as the concentration corresponding to a signal 2.5 standard deviations above the blank. Sample concentrations were back-calculated from the standard curve and reported in pg/mL.

### Western blotting

Cells were washed with DPBS, pelleted, and stored at −80 °C. Pellets were lysed in RIPA buffer (Thermo Fisher Scientific, 89900) supplemented with Halt protease inhibitor (Thermo Fisher Scientific, 78441) and DNase (Thermo Fisher Scientific, 90083), vortexed on ice for 30 min, and sheared through a 32-gauge needle. Lysates were centrifuged at 12,000 rpm for 15 min at 4 °C, and protein concentration was determined by the BCA assay (Pierce, Thermo Fisher Scientific, 23225). Protein (10–20 μg) was denatured in 4× LDS sample buffer (Thermo Fisher Scientific, NP0007) with 2-mercaptoethanol (Gibco, 21985-023) at 100 °C for 5 min, resolved on NuPAGE 4–12% Bis-Tris gels (Thermo Fisher Scientific, NP0322BOX) in 1× MOPS buffer (Thermo Fisher Scientific, NP0001; 80 V for 10 min, then 125 V for 120 min, on ice), and transferred to PVDF using an iBlot 2 system (P0 program: 20 V for 1 min, 23 V for 4 min, 25 V for 2 min).

Membranes were blocked in 5% normal goat serum in 1× TBS-T (Thermo Fisher Scientific, 28360) and incubated overnight at 4 °C with primary antibodies listed in Supplementary Table 2. After three 10-min washes in TBS-T, membranes were incubated with IRDye anti-mouse (LI-COR, 926-80010) or anti-rabbit (LI-COR, 926-80011) secondary antibodies (1:20,000 in blocking buffer, 1 h, RT). Following three further TBS-T washes, membranes were incubated with SuperSignal West Femto substrate (Thermo Fisher Scientific, 34095) for 5 min and imaged on a LI-COR Odyssey system. Densitometric analysis was performed in ImageJ.

### Long-read RNA-seq and qRT–PCR

A total of 5 × 106 TARDBP WT or KO CUTS MNs were plated onto distinct poly-L-ornithine/laminin-coated 10-cm dishes for each technical replicate. After 5 days, cells were dissociated with Accutase, and total RNA was isolated using the RNeasy Mini Kit (Qiagen, 74104) according to the manufacturer’s instructions. For long-read RNA sequencing, 2 μg total RNA was submitted for Iso-Seq library preparation and sequencing (Novogene). For qRT–PCR, 1 μg RNA was reverse transcribed using the High-Capacity cDNA Reverse Transcription Kit with RNase Inhibitor (Thermo Fisher Scientific, 4374966), and cDNA was diluted in nuclease-free water. Reactions were performed in 10 μl volumes in 384-well plates using 200 nM forward and reverse primers, 1.5 ng cDNA, and Fast SYBR Green Master Mix (Thermo Fisher Scientific, 4385612) on a QuantStudio 7 instrument. Primer sequences are listed in Supplementary Table 3. Relative transcript abundance was calculated using the ΔΔCt method, normalized to GAPDH and to a WT control line or DMSO-treated group.

### Long-read RNA-Seq bioinformatics analysis

To generate a transcriptome representing the isoforms present in our samples, we used the grouped, segmented BAM file together with the individual clustered BAM files for each sample, which were provided by Azenta to ALS TDI. Primer sequences and spurious false-positive signals were removed from segmented reads using lima (v2.13.0). Reads were then refined by poly(A) trimming with IsoSeq refine (v4.3.0) to generate full-length non-chimeric (FLNC) reads. FLNC reads were subsequently clustered and error-corrected using IsoSeq Cluster2 (v4.3.0), retaining isoforms supported by at least two FLNC reads. The resulting reads were aligned to the human reference transcriptome (GRCh38) using pbmm2 (v1.16.0). TAMA (v2021_11_03) was then applied to the mapped BAM files to collapse redundant transcript models and remove models arising from fragmented reads or supported by only a single read. This produced a concatenated FASTA file containing all isoforms detected in this study together with previously annotated GRCh38 isoforms. Clustered reads from each sample were then aligned to the assembled transcriptome using pbmm2 (v1.16.0) and quantified with Salmon (v1.10.3).

Differential gene expression (DGE) analysis was performed using edgeR (v4.4.2). Lowly expressed genes were excluded by filtering out genes with fewer than five counts in any sample. Differentially expressed genes (adjusted *P* ≤ 0.05) were subsequently subjected to gene ontology analysis using the enrichWP function in clusterProfiler (v4.14.4).

Differential transcript usage (DTU) analysis was performed using DRIMSeq (v1.34.0). Transcripts with fewer than two counts and less than 10% relative abundance in at least two samples were excluded, and genes with fewer than ten total counts were also removed. This resulted in 8,224 genes and 32,011 isoforms for DTU analysis. All software tools, versions, references, and sources used in this study are listed in Supplementary Table 4.

### Lysosome transport analysis

Lysosome-like vesicle (LV) motility was assessed by live-cell imaging using LysoView vital dyes (Biotium), which accumulate in acidic pH organelles. Day 44 post-plating MNs were labeled with LysoView at a final concentration of 1:1,000 in culture media 24 hours prior to imaging. Time-lapse images were acquired on a BioTek Cytation imager using Gen5 software, capturing one frame per second for 3 minutes per field (181 frames total). Raw images were pre-processed in ImageJ: converted to 8-bit, reordered from Z-stack to time series, and subjected to rolling-ball background subtraction (radius = 50 pixels). Transport density was quantified as a measure of lysosome motility using a custom Jython pipeline implemented in Fiji (ImageJ). Pre-processed 8-bit time-lapse images (181 frames) were binarized by applying a manual intensity threshold and, optionally, cropped to a defined region of interest to exclude image areas lacking cells. For each image, two binary representations were generated: a single-frame image captured at time point zero (*T_0_*), representing the spatial footprint of lysosomes at baseline, and a maximum intensity projection (MIP) collapsed across all frames, representing the cumulative spatial area swept by lysosomes over the entire acquisition. The total thresholded area (µm²) in each representation was quantified using the Analyze Particles function in ImageJ. Transport density was then calculated as the ratio of total MIP area to total *T_0_* area (MIP area / *T_0_* area). Because a stationary lysosome contributes equally to both images, a ratio near 1 indicates minimal movement, while values greater than 1 reflect net displacement of lysosomes across frames.

Per-well results from the MIP and *T_0_* particle analyses were merged using a companion Python script and exported to a single CSV for downstream statistical analysis.

### Compound preparation and treatment

Bortezomib (BTZ; MedChemExpress, HY-10227), MG132 (MedChemExpress, HY-13259), digoxin (Sigma-Aldrich, D6003-100MG), and ouabain (Fisher Scientific, AC161732500) were prepared as DMSO stock solutions and diluted immediately before use in motor neuron plating medium lacking ROCK inhibitor and FUdR.

For acute proteasome inhibition, cryopreserved day 19 CUTS TDP-43 WT MNs were treated at 7 days post-thaw with BTZ (1.11–200 nM) or MG132 (5.62–1000 nM) for 24 h. For cardiac glycoside co-treatment, CUTS TDP-43 WT MNs were treated at 8 days post-thaw with 30 nM BTZ alone or in combination with digoxin or ouabain (0.06–10.00 μM) for 48 h. Vehicle-control wells received matched final concentrations of DMSO (≤0.1%), and cells were fixed at the timepoints indicated in the corresponding figure legends and processed for immunocytochemical analysis as described below..

### Immunocytochemistry

Cryopreserved day 19 MNs were thawed, plated at 10,000 live cells/well in 96-well plates, and fixed in 4% paraformaldehyde at the timepoint indicated in the respective figure legend. After three PBS washes, cells were permeabilized (0.1% Triton X-100 in TBS-T, 5 min), blocked (10% normal donkey serum (NDS) in TBS-T, 1 h), and incubated overnight at 4 °C with primary antibodies diluted in 1% NDS/TBS-T (Supplementary Table 1). Following five TBS-T rinses, cells were incubated for 1 h in secondary antibodies (Supplementary Table 1) in 1% NDS/TBS-T, then counterstained with DAPI (1 μg/mL; Thermo Fisher Scientific, 62248). Image acquisition and quantitative analysis are described below.

### Image acquisition and analysis

For live-cell imaging, cryopreserved day 19 CUTS TDP-43 WT and KO MNs were thawed and plated at 10,000 live cells per well in 96-well plates. At 7 d post-thaw, cells were incubated for 1 h at 37 °C in motor neuron plating medium supplemented with TubulinTracker Deep Red (Thermo Fisher Scientific, T34076; 1:4,000) and Hoechst 33342 (Thermo Fisher Scientific, H1399; 1:500,000).

For immunocytochemistry-based analyses, cryopreserved day 19 MNs were thawed, plated at 10,000 live cells per well in 96-well plates, and fixed at the indicated time points in the respective figure legends. Images were acquired on a Cytation 10 confocal microscope (BioTek) using a 20× objective and a 60-µm spinning disk, and z-stacks were processed as maximum-intensity projections in GEN5 with rolling-disk background subtraction. Quantified parameters are indicated in the respective figure legends. Nuclei were segmented using the DAPI channel, and live nuclei were identified by size-and intensity-based thresholding criteria. All quantitative image-based parameters were measured within the live nuclei population, and all comparison groups were imaged and analyzed using identical acquisition settings and analysis parameters.

Quantitative image analyses were performed in Signals Image Artist (Revvity), and statistical analyses were performed in GraphPad Prism.

### Statistical analysis

All statistical analyses were performed using GraphPad Prism (v10.6.1). Normality of residuals was assessed by visual inspection of Q–Q plots, and homogeneity of variance was evaluated using the Brown–Forsythe test. For comparisons between two groups, equality of variance was assessed using an *F* test, and unpaired, two-tailed Student’s *t*-tests were used for normally distributed datasets with equal variances; when variances were unequal, Welch’s *t*-tests were applied. For comparisons involving more than two groups, datasets meeting assumptions of approximate normality were analyzed using one-way ANOVA followed by Dunnett’s multiple-comparisons test. Datasets deviating from normality were log-transformed and analyzed using Prism’s log-normal one-way ANOVA with Dunnett’s multiple-comparisons test. If transformed data remained non-normal, a nonparametric Kruskal–Wallis test followed by Dunn’s multiple-comparisons test was used. Repeated-measures datasets were analyzed using repeated-measures two-way ANOVA, with time and groups as factors, followed by Dunnett’s multiple-comparisons test; Geisser-Greenhouse correction was applied where sphericity was not assumed. IC50 values were determined by fitting dose–response curves using a four-parameter logistic nonlinear regression model with variable slope. The statistical test used for each dataset, together with the corresponding test statistic, degrees of freedom, and exact *P* value, is reported in the relevant figure legend. Sample sizes (*n*) are defined in the respective figure legends and represent technical replicates unless otherwise stated. Error bars represent mean ± SEM. A significance threshold of *P* < 0.05 was used for all analyses, with significance denoted as follows: \**P* < 0.05, \*\**P* < 0.01, \*\*\**P* < 0.001, \*\*\*\**P* < 0.0001.

## Acknowledgements

This work was generously supported by Augie’s Quest, The Jim Heller Fund, and ALS United Mid-Atlantic. We thank the members of the Cell Biology group at the ALS Therapy Development Institute for their helpful insights and discussion. We thank Alyssa Coyne for kindly sharing qRT-PCR primer sequences. We also thank Target ALS and the Columbia Stem Cell Core, including Dr. Barbara Corneo, Ph.D., Dr. Hynek Wichterle, Ph.D., and Alejandro Garcia-Diaz, for providing the NCRM1-TDT VAChT-iCRE iPSC line and Dr. John Zuris, Ph.D., for kindly sharing the SLEEK donor plasmids. Finally, we thank people with ALS and their families and friends for their support, particularly those who have contributed samples and data to the ALS Research Collaborative.

## Author information

These authors contributed equally: Swetha Gurumurthy, Anushka Bhargava.

## Authors and Affiliations

ALS Therapy Development Institute, Watertown, MA 02472, USA

Swetha Gurumurthy, Anushka Bhargava, Nguyen PT Huynh, Thomas J. Krzystek, Fernando G. Vieira, and Kyle R. Denton.

## Author contribution

A.B. and K.R.D. conceived the project and designed the initial study. Following project expansion, A.B., S.G., and K.R.D. developed the experimental strategy. A.B. and K.R.D. generated and performed initial characterization of the *TARDBP* knockout iPSC lines. A.B. performed qRT-PCR, western blotting, and marker immunocytochemistry to validate *TARDBP* edits, and analyzed TDP-43 loss-of-function phenotypes in knockout iPSC-derived motor neurons. A.B. and T.J.K. performed spot culture, live-cell imaging, and lysosome transport analyses. S.G. designed and performed immunocytochemistry for small-molecule experiments with bortezomib, MG132, digoxin, and ouabain, analyzed CUTS-TDP-43 loss-of-function targets (TDP-43, STMN2, and G3BP1), performed CUTS panel marker staining, and performed TubulinTracker staining. N.P.T.H. performed long-read RNA sequencing and related analyses, and contributed to the corresponding Methods and Results sections. K.R.D. performed qRT-PCR analyses for the bortezomib and digoxin experiments and supervised the study. S.G. and K.R.D. prepared the figures and wrote the original draft of the manuscript. S.G., A.B., T.J.K., N.P.T.H., F.G.V., and K.R.D. critically revised the manuscript. All authors approved the final manuscript.

## Data availability

The data that support the findings of this study are available from the corresponding author upon reasonable request. The Source Data file containing the data underlying all graphs and figures will be provided to the editors and reviewers for peer review if requested and will be made available with the published article.

## Code availability

Publicly available scripts, pipelines, and software tools used in this study are listed in Supplementary Table 4, together with version information, references, and source details. Any study-specific analysis parameters are available from the corresponding author upon reasonable request.

## Supplementary Tables

**Supplementary Table 1.**
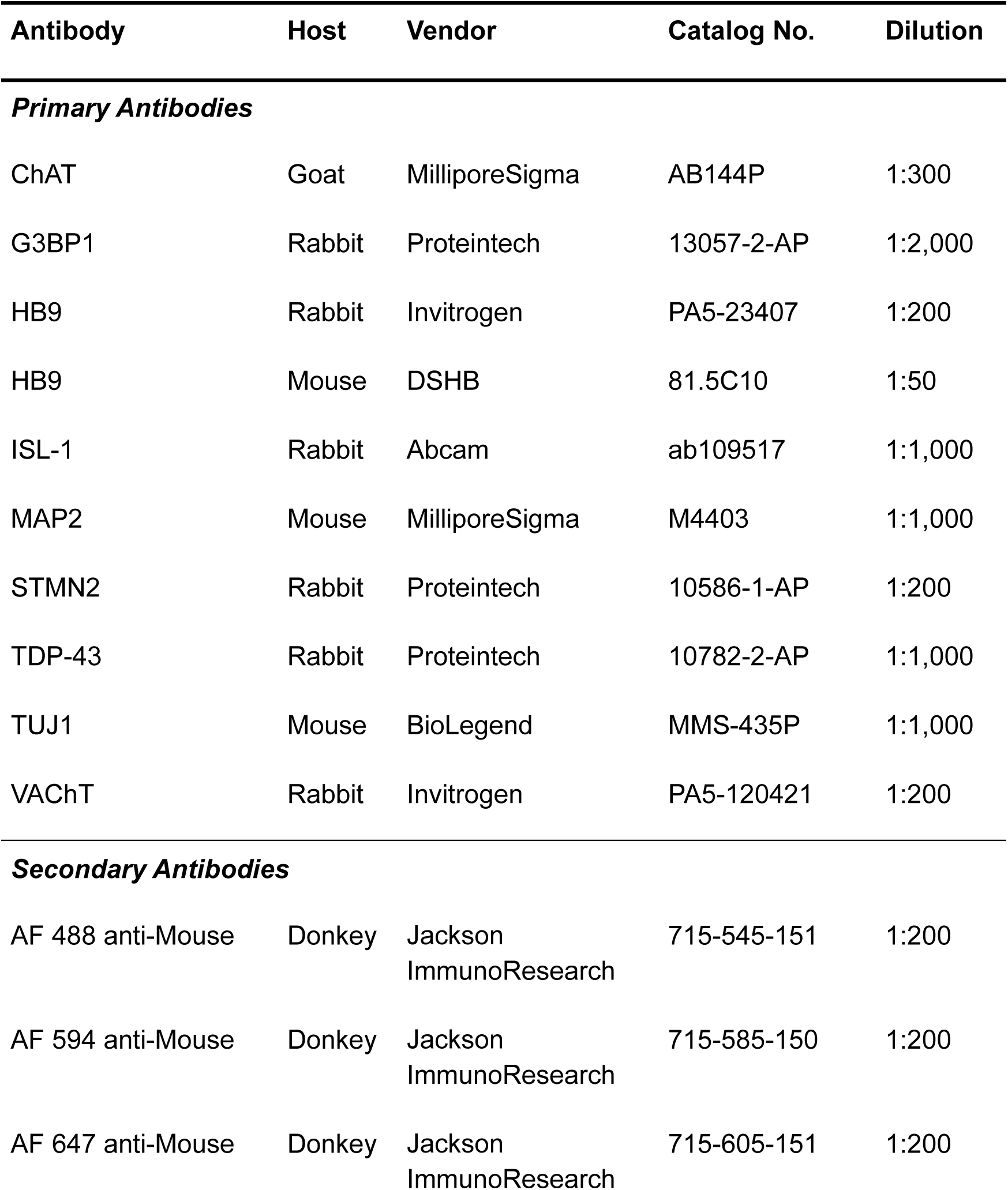
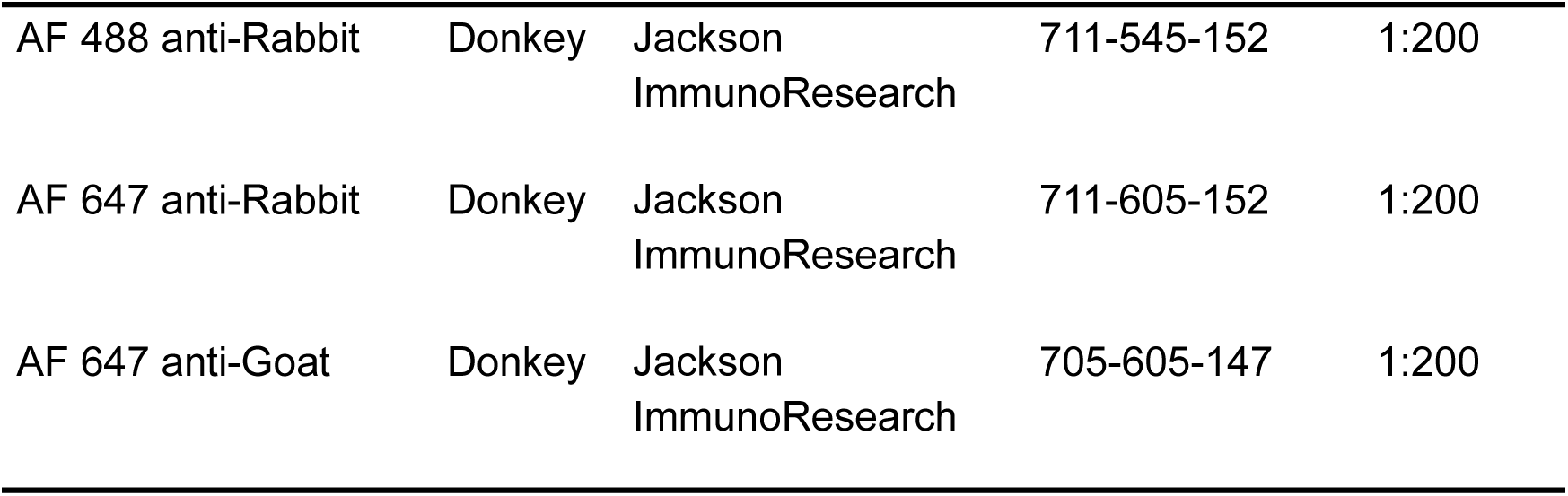
Primary and secondary antibodies used for immunocytochemistry.

**Supplementary Table 2.** Primary antibodies used for western blotting.

| **Antibody** | **Host** | **Vendor** | **Catalog no.** | **Dilution** |
| --- | --- | --- | --- | --- |
| ChAT | Goat | MilliporeSigma | AB144P | 1:500 |
| GAPDH | Mouse | Invitrogen | AM4300 | 1:10,000 |
| MAP2 | Mouse | MilliporeSigma | M4403 | 1:1,000 |
| STMN2 | Rabbit | Proteintech | 10586-1-AP | 1:2,000 |
| TDP-43 | Rabbit | Proteintech | 10782-2-AP | 1:1,000 |
| UNC13A | Rabbit | Synaptic Systems | 126-103 | 1:1,000 |
| α-Tubulin | Mouse | MilliporeSigma | T8203 | 1:20,000 |

**Supplementary Table 3.**
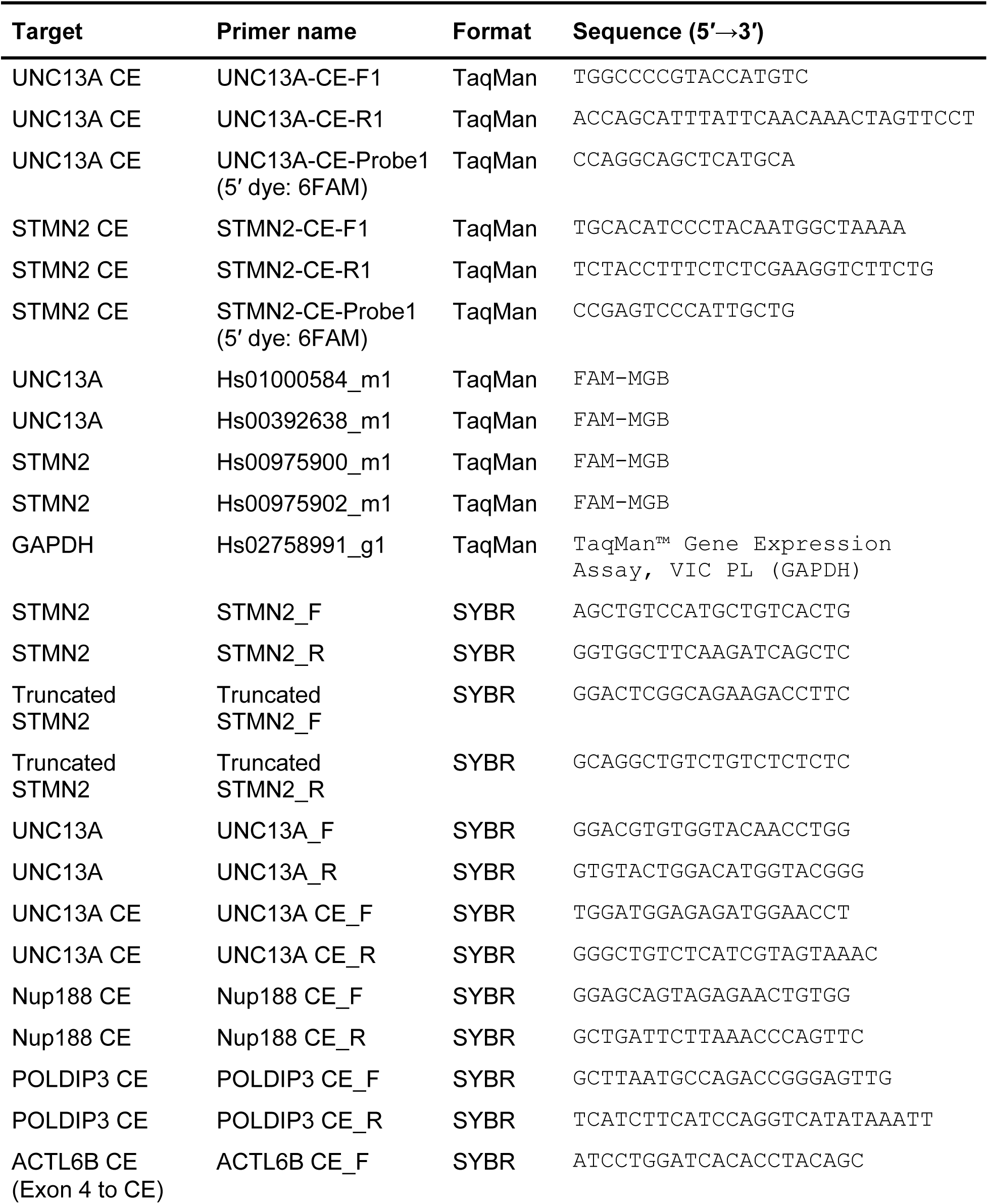
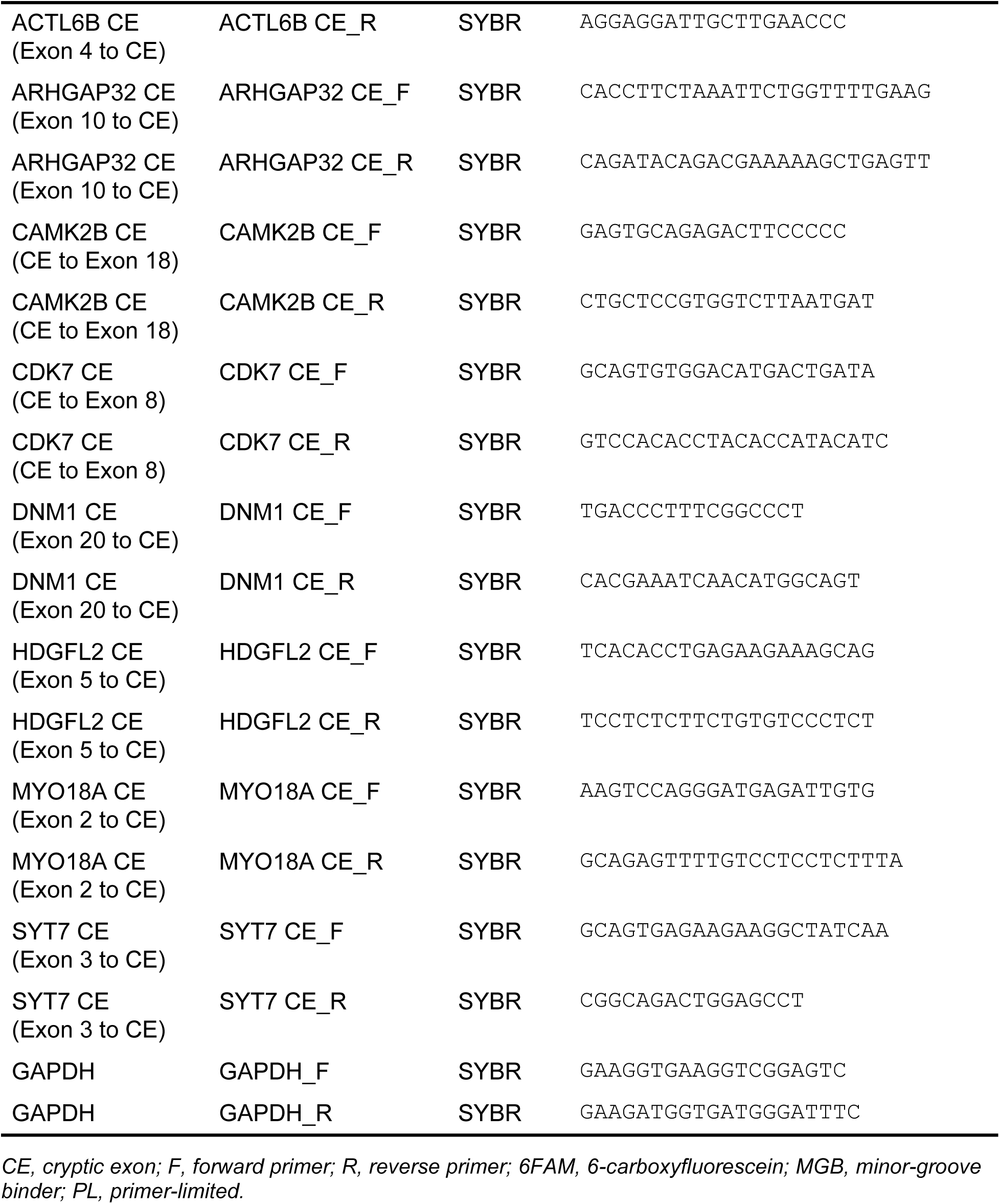
qRT-PCR primers and probes.

**Supplementary Table 4.** Software and tools used in this study.

| Software | Version | Reference | Source |
| --- | --- | --- | --- |
| lima | 2.13.0 | — | https://github.com/PacificBiosciences/barcoding |
| IsoSeq | 4.3.0 | — | https://isoseq.how/ |
| pbmm2 | 26.1.0 | — | https://github.com/PacificBiosciences/pbmm2 |
| TAMA | 2021_11_03 | 30 | https://github.com/GenomeRIK/tama/ |
| salmon | 1.10.3 | 31 | https://combine-lab.github.io/salmon/ |
| DRIMSeq | 1.34.0 | 32 | https://github.com/gosianow/DRIMSeq |
| edgeR | 4.4.2 | 33 | https://bioconductor.org/packages/release/bioc/html/edgeR.html |
| clusterProfiler | 4.14.4 | 34,35 | https://bioconductor.org/packages/release/bioc/html/clusterProfiler.html |

*Reference numbers correspond to the main manuscript bibliography. Tools without associated publications are indicated by an em-dash; see the Source column for repository URLs.*

**Supplementary Table 5.** Motor neuron yield and fold expansion.

| Cell line | Line code | iPSCs plated (day -1) | Total MNs (day 19) | Fold expansion (day 19 MNs / day -1 iPSCs) |
| --- | --- | --- | --- | --- |
| CUTS TDP-43 WT | 42 | $1.701 \times 10^6$ | $7.38 \times 10^8$ | 433.9 |
| CUTS TDP-43 KO | 47 | $1.701 \times 10^6$ | $4.72 \times 10^7$ | 27.7 |
| CUTS TDP-43 KO | 50 | $1.701 \times 10^6$ | $4.58 \times 10^7$ | 26.9 |

*WT, wild-type; KO, knockout. Fold expansion is calculated as the ratio of total live MNs recovered at day 19 to live iPSCs plated at day -1.*

## Supplementary Figures

**Supplementary Figure 1:**
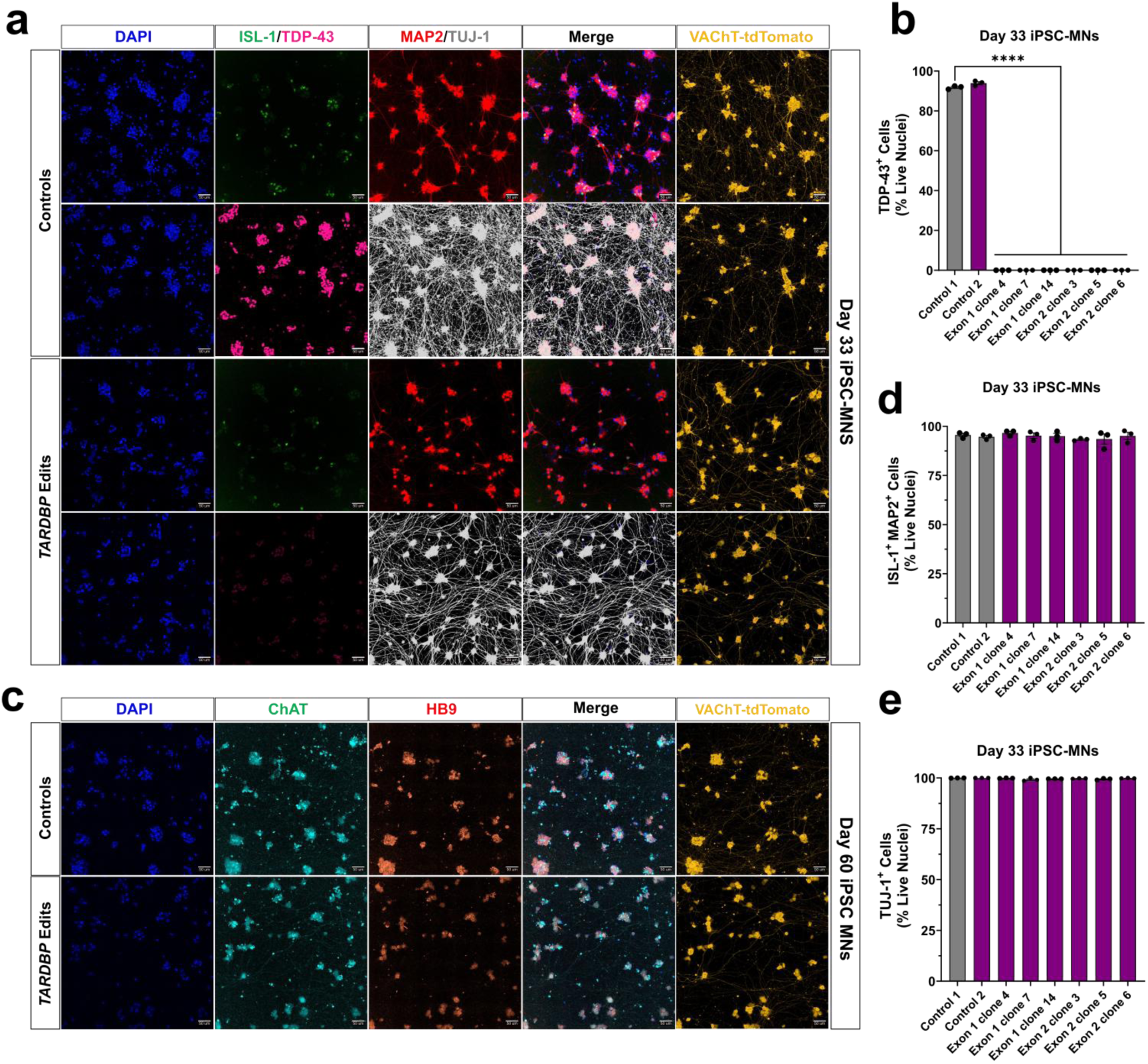
Characterization of neuronal marker expression in *TARDBP* knockout iPSC-derived motor neurons. **(a)** Representative immunofluorescence images of day 33 iPSC-MNs showing DAPI, ISL-1, TDP-43, MAP2, TUJ-1, Merge, and VAChT-tdTomato. tdTomato expression is driven by the VAChT promoter. **(b)** Quantification of TDP-43^+^ cells (% Live Nuclei), defined by TDP-43 intensity above the no-primary-control threshold, across isogenic control and *TARDBP* KO clones at day 33. One-way ANOVA with Dunnett’s test vs Control 1; *F*(7, 16) = 19929, *P* < 0.0001. **(c)** Representative immunofluorescence images of day 60 iPSC-MNs showing DAPI, ChAT, HB9, merged signal, and VAChT-tdTomato. **(d)** Quantification of ISL-1^+^ MAP2^+^ cells (% Live Nuclei) at day 33. **(e)** Quantification of TUJ-1^+^ cells (% Live Nuclei) at day 33. **(a, c)** Scale bars, 50 µm. **(b, d, e)** Bars represent mean ± SEM; *n* = 3 technical replicates per condition. \*\*\*\**P* < 0.0001.

**Supplementary Figure 2.**
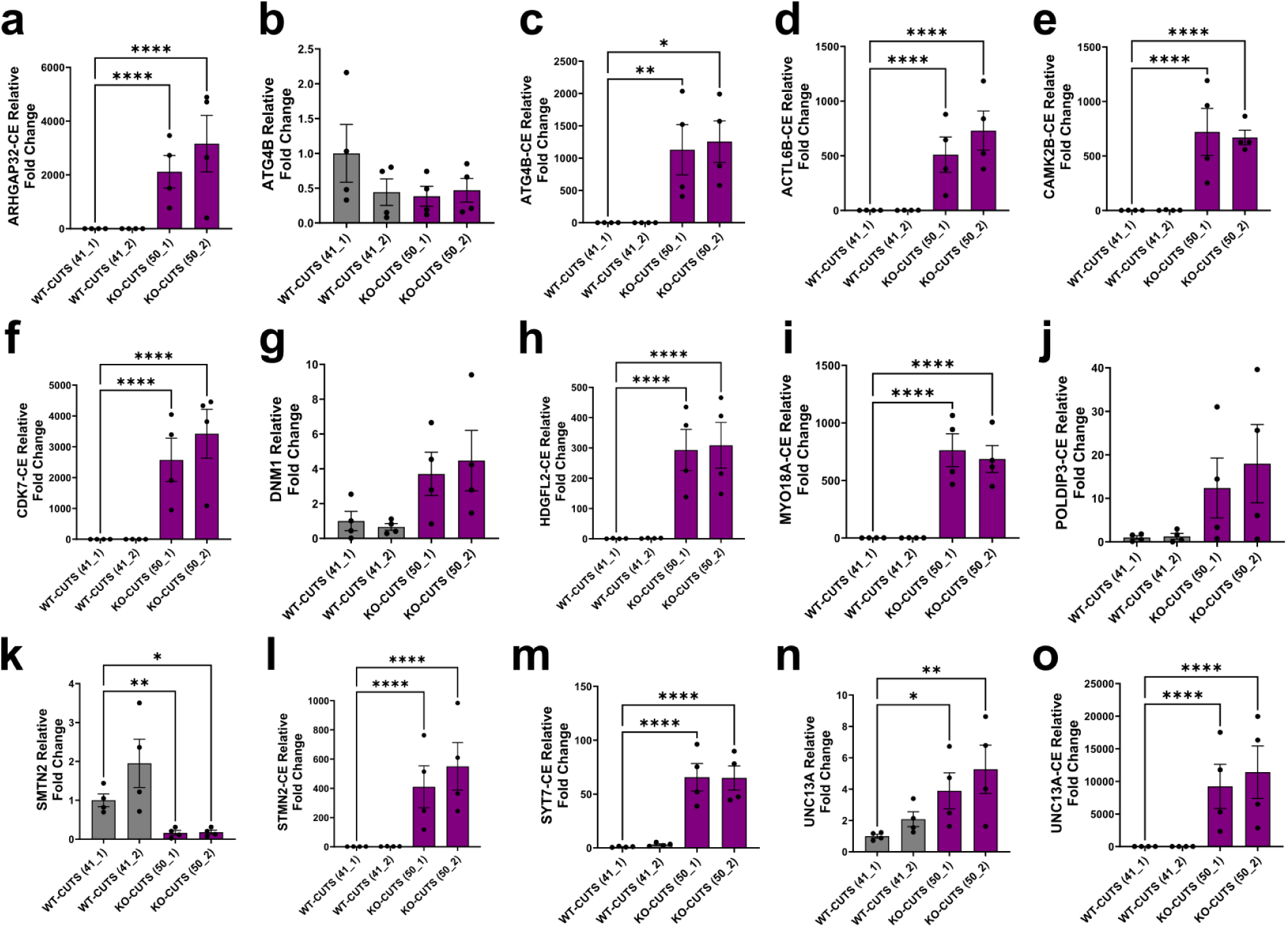
qRT-PCR validation of TDP-43-dependent cryptic exon expression in CUTS TDP-43 wild-type and knockout motor neurons. **(a–o)** Quantification of relative fold change by qRT-PCR in CUTS TDP-43 WT (lines 41_1 and 41_2) and KO (lines 50_1 and 50_2) MNs. **(a)** ARHGAP32-CE (*F*(3,12) = 119.8, *P* < 0.0001). **(b)** ATG4B (*F*(3,12) = 1.271, *P* = 0.3285). **(c)** ATG4B-CE (*F*(3.000, 6.424)= 43.06, *P* = 0.0002). **(d)** ACTL6B-CE (*F*(3,12) = 109.1, *P* < 0.0001). **(e)** CAMK2B-CE (*F*(3,12) = 33.57, *P* < 0.0001). **(f)** CDK7-CE (*F*(3,12) = 102.0, *P* < 0.0001). **(g)** DNM1-CE (*F*(3,12) = 2.983, *P* = 0.0738). **(h)** HDGFL2-CE (*F*(3,12) = 113.8, *P* < 0.0001). **(i)** MYO18A-CE (*F*(3,12) = 322.4, *P* < 0.0001). **(j)** POLDIP3-CE (*F*(3,12) = 2.393, *P* = 0.1194). **(k)** STMN2 (*F*(3,12) = 12.17, *P* = 0.0006). **(l)** STMN2-CE (*F*(3,12) = 133.6, *P* < 0.0001). **(m)** SYT7-CE (*F*(3,12) = 94.03, *P* < 0.0001). **(n)** UNC13A (*F*(3,12) = 5.921, *P* = 0.0102). **(o)** UNC13A-CE (*F*(3,12) = 62.22, *P* < 0.0001). **(a, d–o)** One-way ANOVA on log-transformed values with Dunnett’s test vs WT-CUTS (41_1). **(b, g)** Ordinary one-way ANOVA with Dunnett’s test vs WT-CUTS (41_1). **(c)** Brown-Forsythe ANOVA on log-transformed values with Dunnett’s test vs WT-CUTS (41_1). **(a–o)** Bars represent mean ± SEM; *n* = 4 technical replicates per condition. \**P* < 0.05, \*\**P* < 0.01, \*\*\**P* < 0.001, \*\*\*\**P* < 0.0001.

**Supplementary Figure 3.**
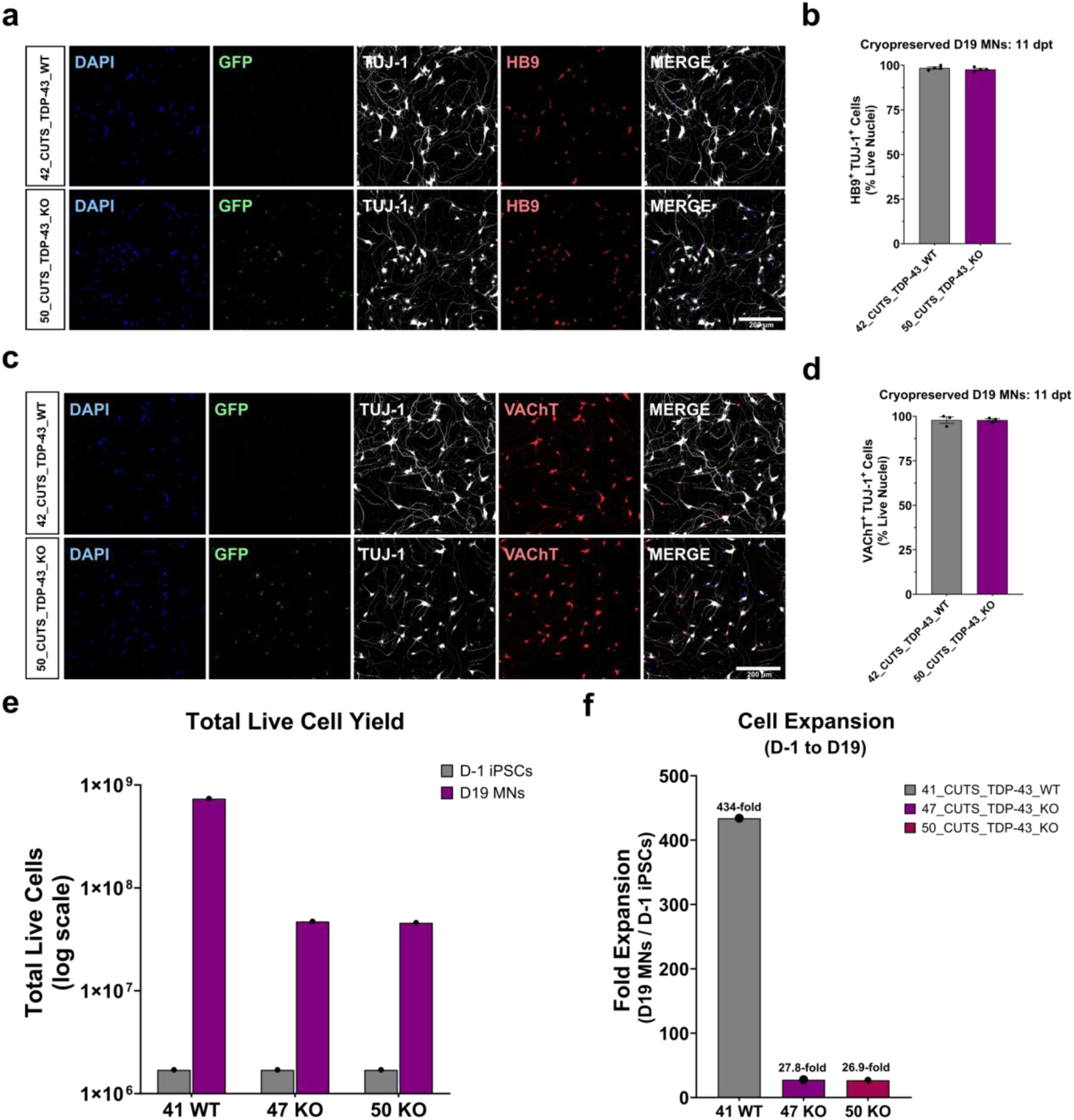
Characterization of CUTS TDP-43 WT and KO motor neuron identity and differentiation yield**. (a)** Representative immunofluorescence images of cryopreserved D19 CUTS TDP-43 WT (line 42) and TDP-43 KO (line 50) MNs, fixed at 11 days post-thaw, showing DAPI, cryptic GFP, TUJ-1, and HB9. **(b)** Quantification of HB9^+^ TUJ-1^+^ cells (% Live Nuclei) in WT and KO MNs (*n* = 4 technical replicates per condition). **(c)** Representative immunofluorescence images of cryopreserved D19 WT (line 42) and KO (line 50) MNs, fixed at 11 days post-thaw, highlighting DAPI, cryptic GFP, TUJ-1, and VAChT. **(d)** Quantification of VAChT^+^ TUJ-1^+^ cells (% Live Nuclei) in WT and KO MNs (*n* = 3 technical replicates per condition). **(e)** Total live cell differentiation yield (log scale) at D−1 (iPSCs) and D19 (MNs) for CUTS TDP-43 WT (line 41) and KO (lines 47, 50). **(f)** Fold expansion from D-1 iPSCs to D19 MNs for CUTS TDP-43 WT (line 41; 434-fold), CUTS TDP-43 KO (line 47; 27.8-fold), and CUTS TDP-43 KO (line 50; 26.9-fold). **(a, c)** Scale bars, 200 μm. **(b, d)** Bars represent mean ± SEM. \**P* < 0.05, \*\**P* < 0.01, \*\*\**P* < 0.001, \*\*\*\**P* < 0.0001.

**Supplementary Figure 4.**
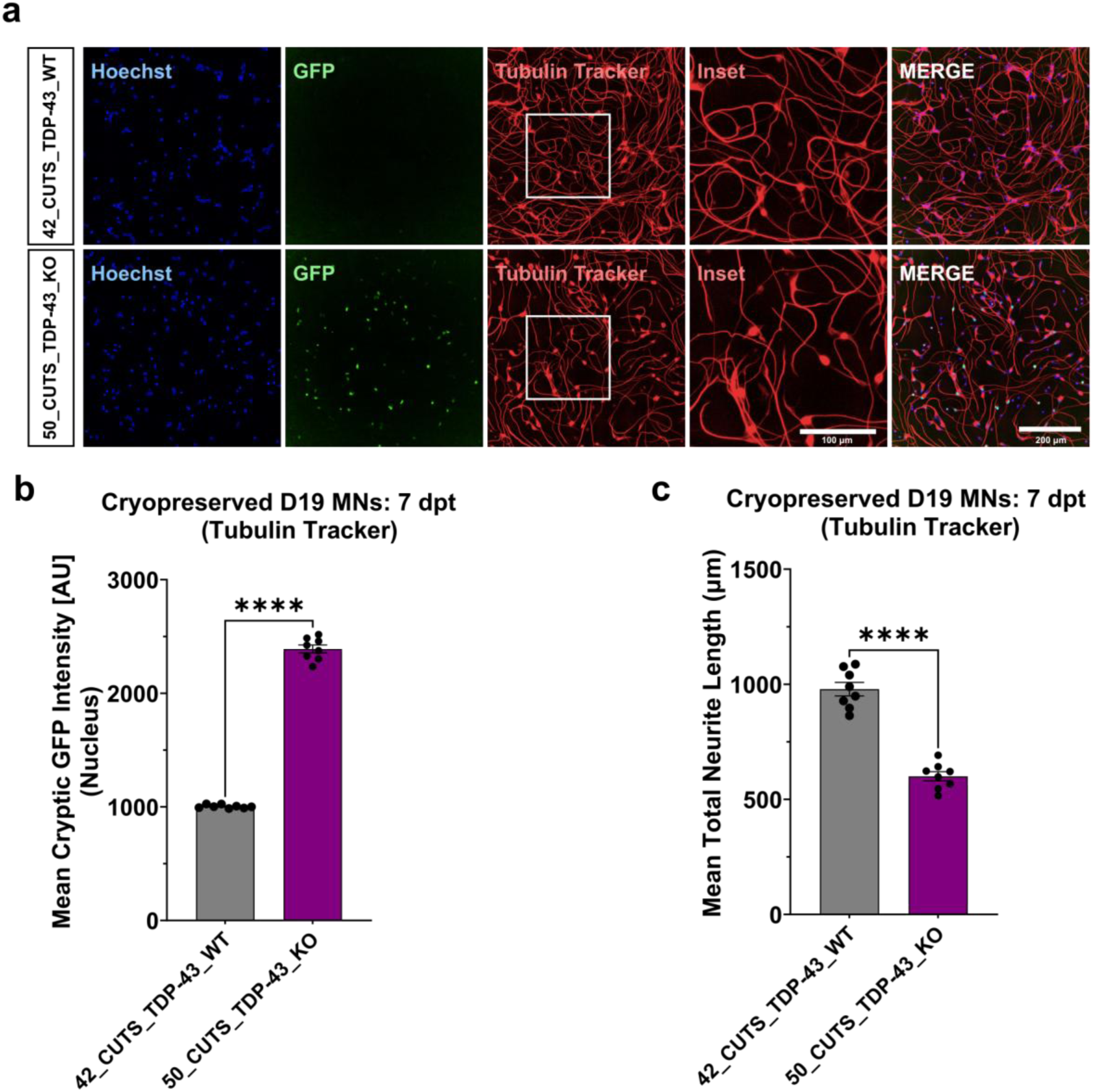
CUTS TDP-43 knockout motor neurons exhibit reduced neurite outgrowth and increased cryptic GFP reporter expression. **(a)** Representative confocal images of live cryopreserved day 19 CUTS TDP-43 WT (line 42) and TDP-43 KO (line 50) MNs at 7 days post-thaw, showing Hoechst, cryptic GFP, and TubulinTracker Deep Red and merge. Insets show a cropped view of the indicated TubulinTracker region. Scale bar, 200 μm; inset, 100 μm. **(b)** Quantification of mean nuclear cryptic GFP intensity in WT and KO MNs (two-tailed Welch’s unpaired *t*-test; *t*(7.331) = 39.51, *P* < 0.0001). **(c)** Quantification of mean total neurite length (μm) in WT and KO MNs (two-tailed unpaired *t*-test; *t*(14) = 10.62, *P* < 0.0001). **(b, c)** Bars represent mean ± SEM; *n* = 8 technical replicates per condition. \**P* < 0.05, \*\**P* < 0.01, \*\*\**P* < 0.001, \*\*\*\**P* < 0.0001.

**Supplementary Figure 5.**
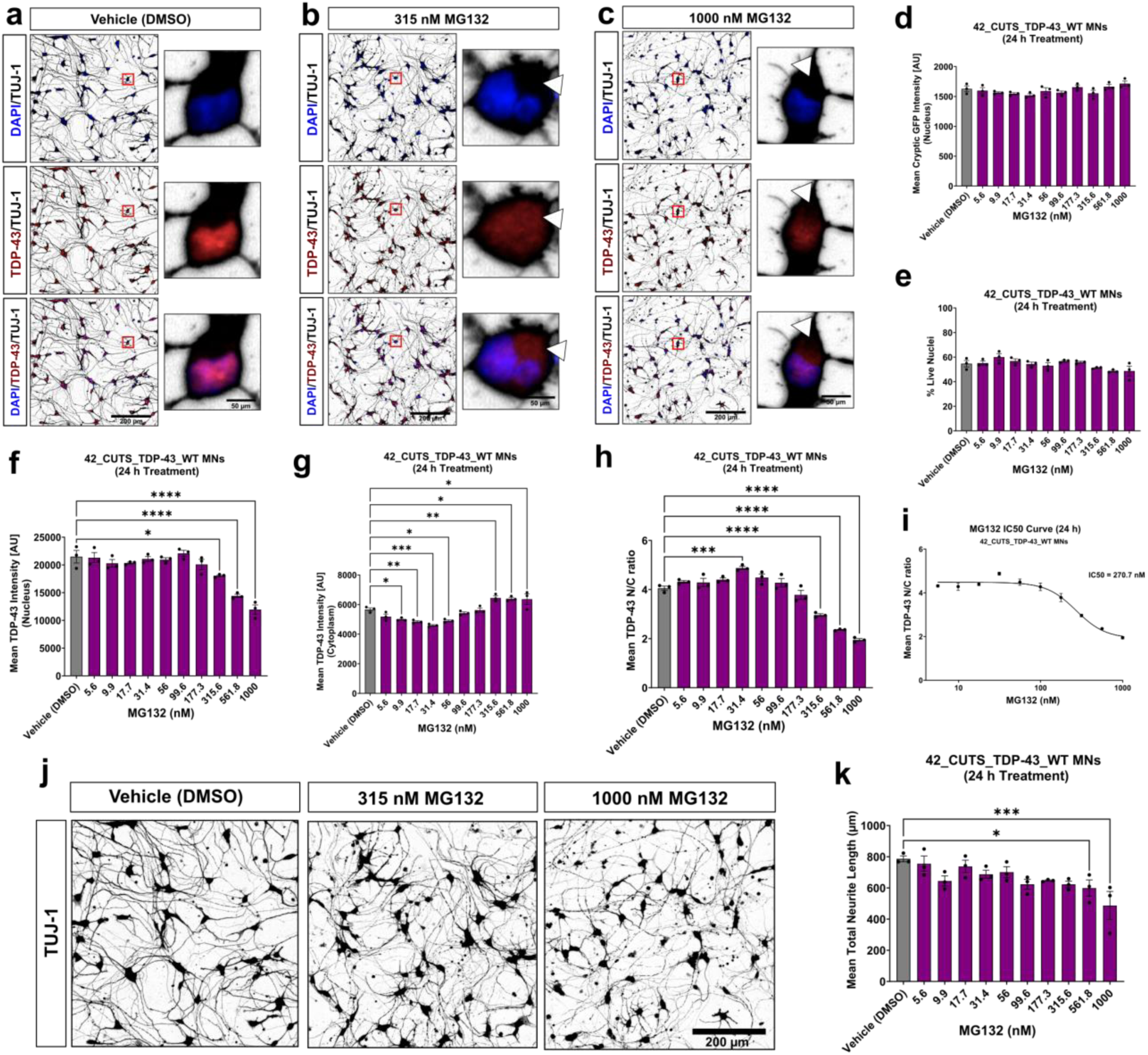
MG132 induces dose-dependent TDP-43 mislocalization in CUTS TDP-43 wild-type motor neurons. **(a–c)** Representative immunofluorescence images of CUTS TDP-43 WT MNs (line 42) treated for 24 h with DMSO vehicle (a), 315 nM MG132 (b), or 1000 nM MG132 (c), showing DAPI/TUJ-1, TDP-43/TUJ-1, and merged channels. Insets highlight subcellular TDP-43 localization; white arrowheads in (b) and (c) indicate cytoplasmic TDP-43 mislocalization. Scale bar, 200 μm; insets, 50 μm. **(d, e)** Quantification of mean nuclear cryptic GFP intensity (d; *F*(10, 22) = 2.245, *P* = 0.0548) and % live nuclei (e; *F*(10, 22) = 2.810, *P* = 0.0208) across MG132 titration (24 h) in CUTS TDP-43 WT MNs (line 42). **(f–h)** Quantification of nuclear TDP-43 intensity (f; *F*(10, 22) = 23.06, *P* < 0.0001), cytoplasmic TDP-43 intensity (g; *F*(10, 22) = 20.19, *P* < 0.0001), and TDP-43 mislocalization ratio (h; *F*(10, 22) = 69.96, *P* < 0.0001) across MG132 titration (24 h). **(i)** IC50 curve for TDP-43 mislocalization (24 h); IC50 = 270.7 nM, fitted using a four-parameter variable-slope model (log[inhibitor] vs. response). **(j)** Representative TUJ-1 immunofluorescence images of WT MNs treated with Vehicle (DMSO), 315 nM MG132, or 1000 nM MG132 for 24 h. Scale bar, 200 μm. **(k)** Mean total neurite length (μm) across MG132 titration (24 h; *F*(10, 22) = 3.895, *P* = 0.0037). **(a–c)** Scale bars, 200 μm; insets, 50 μm. **(d–h, k)** Bars represent mean ± SEM; *n* = 3 technical replicates per condition. One-way ANOVA with Dunnett’s test vs Vehicle (DMSO). \**P* < 0.05, \*\**P* < 0.01, \*\*\**P* < 0.001, \*\*\*\**P* < 0.0001.

